# Spatiotemporal Single-Cell Roadmap of Human Skin Wound Healing

**DOI:** 10.1101/2024.08.30.609923

**Authors:** Zhuang Liu, Dongqing Li, Xiaowei Bian, Lihua Luo, Åsa K. Björklund, Li Li, Letian Zhang, Yongjian Chen, Lei Guo, Juan Gao, Chunyan Cao, Jiating Wang, Wenjun He, Yunting Xiao, Liping Zhu, Karl Annusver, Huda Gopee, Clare L. Bennett, Maria Kasper, Muzlifah Haniffa, Pehr Sommar, Ning Xu Landén

## Abstract

Wound healing is vital for human health, yet the details of cellular dynamics and coordination in human wound repair remain largely unexplored. To address this, we conducted single-cell and spatial transcriptomics analyses on human skin and acute wound tissues through inflammation, proliferation, and remodeling phases of wound repair from the same individuals, monitoring the cellular and molecular dynamics of human skin wound healing at an unprecedented spatiotemporal resolution. This singular roadmap reveals the cellular architecture of the wound margin and identifies FOSL1 as a critical driver of re-epithelialization. It shows that pro-inflammatory macrophages and fibroblasts sequentially support keratinocyte migration like a relay race across different healing stages. Comparison with single-cell data from venous and diabetic foot ulcers uncovers a link between failed keratinocyte migration and impaired inflammatory response in chronic wounds. Additionally, comparing human and mouse acute wound transcriptomes underscores the indispensable value of this roadmap in bridging basic research with clinical innovations.

**Highlights:**

- A spatiotemporal atlas to explore human *in vivo* gene expression across wound healing
- Wound margin architecture unveils a model for human re-epithelialization
- Distinct healing challenges in venous ulcers and diabetic foot ulcers
- Unique human healing traits emerge in cellular heterogeneity and gene expression

## INTRODUCTION

Wound healing is a vital process for skin integrity, progressing through three overlapping phases: inflammation, proliferation, and remodeling^1,2^. These stages are driven by complex interactions among diverse cell types, such as keratinocytes, fibroblasts, and immune cells. Re-epithelialization, crucial during the proliferation phase, requires the migration and proliferation of epidermal keratinocytes to cover the wound. Failed re-epithelialization is a common issue in chronic wounds like diabetic foot ulcers (DFU) and venous ulcers (VU), impacting millions globally each year^3^. There’s an urgent need for more effective wound therapies, yet progress is hindered by limited knowledge of human skin wound healing. While animal models, mainly rodents, have provided foundational insights, significant anatomical and physiological differences between humans and rodents limit their applicability, often resulting in high clinical trial failure rates for new treatments^4,5^.

Significant efforts have been made to characterize human wound tissues using conventional and advanced techniques such as single-cell RNA sequencing (scRNA-seq) and spatial transcriptomic sequencing (ST-seq). Existing research has primarily analyzed pathological wounds, such as pressure ulcers^6^ and DFUs^7,8^, as well as pathological scars^9–11^, as these samples are more readily obtained during treatment. However, a comprehensive cellular atlas detailing normal skin wound healing in humans over time is still missing. This knowledge gap is crucial for identifying critical cellular and molecular mediators of wound healing, understanding obstacles to healing in chronic wounds, and validating the clinical relevance of findings from animal and *in vitro* models.

In this study, we utilized scRNA-seq and ST-seq to monitor gene expression dynamics at the single-cell level across different healing phases in the same human donors. These efforts led to the development of a comprehensive spatiotemporal cell atlas of human skin wound healing, accessible for interactive exploration online (https://www.xulandenlab.com/tools). Through this atlas, we identified a novel epidermal wound margin structure and fundamental regulatory mechanisms driving re-epithelialization in humans. Our comparative studies with chronic wounds revealed diverse pathological changes specific to their causes and prognoses. Additionally, the comparison of human and mouse acute wound transcriptomes showed both shared and unique healing mechanisms, highlighting this atlas’s crucial role in translating fundamental discoveries into new therapeutic approaches.

## RESULTS

### A spatiotemporal map of human skin wound healing

In our study, we created controlled full-thickness skin wounds on consenting volunteers (**Figure 1a**, **Table S1**). We collected samples from pre-wound intact skin and the entire concentric wound edge at each phase of healing: immediate inflammation (one day post-injury, Wound1), proliferation (seven days post-injury, Wound7), and remodeling (30 days post-injury, Wound30), all from the same individuals. This approach ensures that the data reflect the healing process consistently within individual biological contexts rather than introducing variability by comparing different stages from different donors. We analyzed serial human acute wound samples from three donors using 10x Genomics scRNA-seq and four donors using Visium ST-seq. Biopsies from two donors were analyzed using both techniques. Additionally, we performed scRNA-seq on VU biopsies from four patients, with each biopsy consisting of 50% wound-edge and 50% wound-bed tissues (**Figure 1a**, **Table S1**). We also analyzed five intact skin samples from healthy donors matched by age and gender to the VU patients.

**Figure 1.**
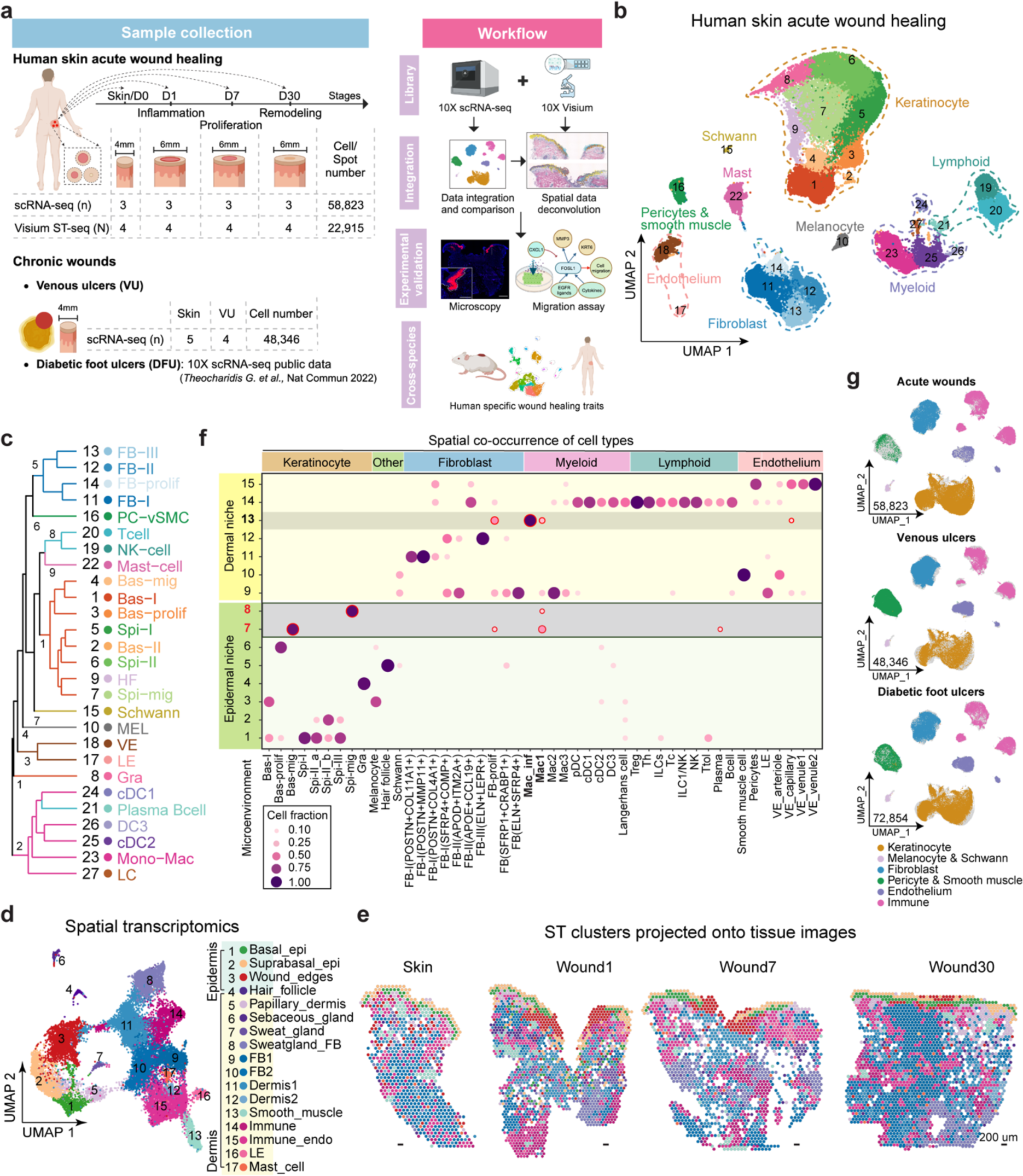
A spatiotemporal map of human skin wound healing. (**a**) Schematic outline of the study. (**b**) Uniform manifold approximation and projection (UMAP) of cell populations in human acute wound scRNA-seq. (**c**) Dendrogram illustrating cell clusters’ relatedness based on a distance matrix constructed in gene expression space. (**d**) UMAP of spot clusters in human acute wound ST-seq. Colors represent different clusters. (**e**) Spatial projection of cell types onto hematoxylin and eosin (H&E) images of human skin and acute wounds. (**f**) Dot plot showing spatial co-occurrence of different cell types using non-negative matrix factorization (NMF) analysis of ST-seq deconvolution results. Dot size and color represent cell fractions normalized across niches for each cell type. (**g**) UMAP of main cell types in integrated human acute and chronic wound scRNA-seq datasets. epi, epidermis; endo, endothelium; Bas-/Spi-/Gra-, basal/spinous/granular keratinocyte; - prolif/-mig, proliferating/migrating cells; HF, hair follicle; MEL, melanocyte; FB, fibroblast; Mono-mac, monocyte and macrophage; Mac_inf: inflammatory macrophage; cDC/pDC, conventional/ plasmacytoid dendritic cell; LC, Langerhans cell; NK-cell, natural killer cell; Th, T helper cell; Treg, regulatory T cell; ILC, innate lymphoid cell; Tc: cytotoxic T cell; Ttol, tolerant T cell; PC-vSMC, pericyte and vascular smooth muscle cell; LE/VE, lymphatic/vascular endothelial cell.

After filtering out potential doublets and low-quality cells using the Scrublet^12^, DoubletFinder^13^, and Seurat packages^14^, we analyzed 58,823 cells from 12 scRNA-seq libraries of human skin and acute wounds at days 1, 7, and 30 from three donors (**Figure S1a-c**). Using graph-based clustering with the Louvain algorithm^14^, we identified 27 cell clusters, which were further grouped into nine main cell types based on differentially expressed genes (DEGs) and known markers^15,16^: keratinocytes (*KRT5*^high^ or *KRT10* ^high^), fibroblasts (*COL1A1^+^*), myeloid cells (*LYZ^+^*, *HLA-DRA^+^*), lymphoid cells (*CD3D^+^* or *NKG7^+^*), endothelial cells (*PECAM1^+^*), mast cells (*TPSAB1^+^*, *TPSB2^+^*), pericytes and smooth muscle cells (*ACTA2^+^*, *MYH11^+^*), melanocytes (*TYRP1^+^*, *PMEL^+^*), and Schwann cells (*SOX10^+^*, *SOX2^+^*) (**Figure 1b, c**, **Figure S1d**, **Table S2**). Integration and comparison with public scRNA-seq data from human adult skin confirmed the accuracy and comprehensiveness of our cell type annotations (**Figure S1e, f**)^16^.

Our ST-seq analysis of human skin and acute wounds identified 22,915 spots expressing over 100 genes (**Figure S2a, b**), organized into 17 clusters (**Figure 1d, e**, **Figure S2c, d**, **Table S2**). The epidermal compartment included basal keratinocytes (*KRT15^+^*, *COL17A1^+^*), suprabasal cells (*KRT1^+^*, *KRT2^+^*, *LORICRIN^+^*), hair follicles (*HR^+^*, *FZD7^+^*, *KRT75^+^*)^17,18^, and a wound-edge cluster (*KRT6A/B/C^+^*, *KRT16^+^*, *KRT17^+^*, *S100A8/9^+^*). The dermal clusters comprised fibroblasts (*ADAM12^+^*, *POSTN^+^*, *MMP2^+^*, *FBLN1^+^*)^19,20^, sweat glands (*KRT77^+^*, *DCD^+^*, *SCGB1D2^+^*, *MUCL1^+^*)^21^, sebaceous glands (*MGS11^+^*, *FADS2^+^*, *KRT79^+^*)^17,18^, immune cells (*FABP4^+^*, *CD36^+^*, *CD163^+^*), endothelial cells (*VWF^+^*, *CD74^+^*, *CCL21^+^*, *LYVE1^+^*), smooth muscle cells (*MYL9^+^*, *TAGLN^+^*)^22^, and a mast cell cluster (*TPSB2^+^*, *MS4A2^+^*).

To map the spatial positions of cells identified by scRNA-seq within their native environments, we applied Cell2location analysis^23^ to our ST-seq data, resulting in a spatiotemporal single-cell atlas of human skin wound healing. Using non-negative matrix factorization^23^, we distinguished the epidermal and dermal niches, revealing specific microenvironments where different cell types co-localized (**Figure 1f**, **Figure S2e**). For instance, the dermal vascular microenvironment (niches 10 and 15) contains smooth muscle cells, pericytes, and various endothelial cells, while the basal epidermal layer (niche 3) comprises basal-I keratinocytes and melanocytes (**Figure S2e**). These microenvironments are crucial for understanding spatially restricted cell-to-cell communication during wound healing. Notably, our analysis highlights interactions between macrophages with migrating keratinocytes (niches 7 and 8) and proliferating fibroblasts (niche 13), underscoring the role of wound inflammation in repair beyond infection defense and debris clearance (**Figure 1f**, **Figure S2e**).

To deepen our understanding of chronic wound pathology, we integrated scRNA-seq data from human acute wounds (skin, wound1, wound7, and wound30) with data from VU (four VUs vs. five skin) and public DFU data^8^ using Harmony^24^ (**Figure 1g**, **Figure S1g-i**). Notably, the DFU study compared foot skin from nine non-diabetic individuals to DFU wound-edge tissues from 11 diabetic patients. These DFU cases were divided into two groups based on 12-week healing outcomes: healed (DFU_H, n=7) and non-healed (DFU_NH, n=4)^8^. This integration allows us to directly compare human acute wounds to chronic wounds of different etiologies and identify pathological barriers that hinder wound closure.

### Spatiotemporal dissection of human wound re-epithelialization

Recognizing that impaired re-epithelialization is a common challenge in pathological wounds^3^, our study aimed to better understand this process in human wounds. Using scRNA-seq, we classified keratinocytes from human skin and acute wounds into nine sub-clusters (**Figure 2a**). These clusters include three basal types: Bas-I cells (*ASS1^+^*, *POSTN^+^*), a proliferating type (Bas-prolif: *STMN1^+^*, *TOP2A^+^*), and a migrating type (Bas-mig) expressing matrix metalloproteinases (*MMPs*) and *FGFBP1*. We also identified five spinous (Spi) types, ranging from Spi-I/IIa/b with metallothionein expression to Spi-III with immune response genes (*TNFSF10, IRF, CCL27*), and a migrating spinous cluster (Spi-mig) expressing *KRT6* genes and S100 proteins. The granular cluster (Gra) displays late differentiation markers (e.g., *FLG, LORICRIN*) (**Figure 2b**, **Table S3**). Analysis of cell proportions and graph-based differential abundance testing using Milo^25^ revealed a notable early increase in migrating keratinocytes (Bas-mig and Spi-mig) during the initial inflammatory phase of wound healing (Wound1). This increase is contrasted by a decrease in differentiated keratinocytes, such as Spi-II and Gra, compared to normal skin (**Figure 2c, d**, **Figure S3a**). These observations were supported by deconvolution analysis of our prior bulk RNA-seq data from human skin and wound samples using the same *in vivo* human wound healing model^26^ (**Figure 2e**, **Figure S3b**). Additionally, we noted increased proliferating keratinocytes at Wound7, suggesting a robust proliferative response to facilitate wound healing (**Figure 2c**, **Figure S3c**).

**Figure 2.**
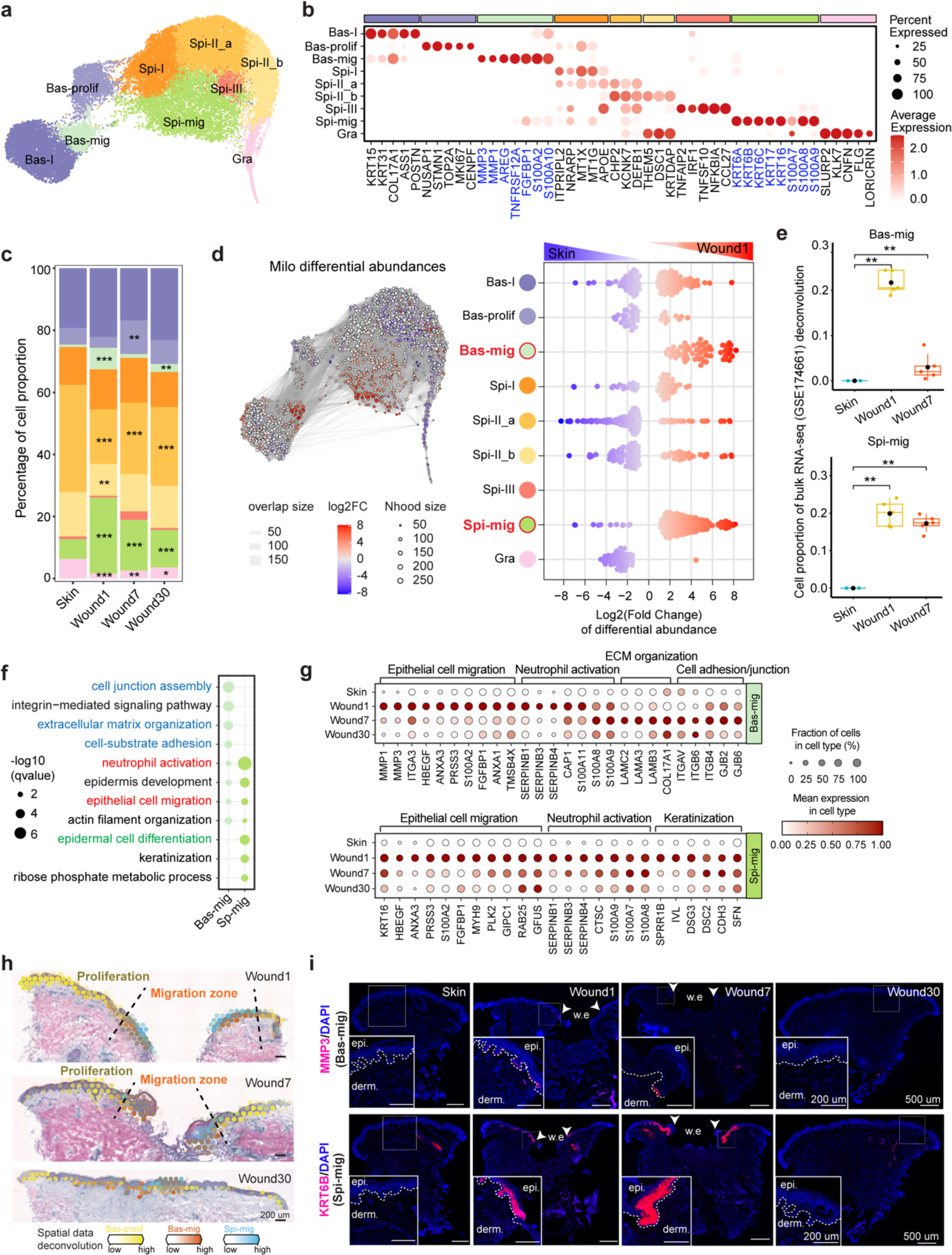
Spatiotemporal dissection of human wound re-epithelialization. (**a**) UMAP of keratinocyte subclusters in human acute wound healing. (**b**) Dot plot showing the expression of each cluster’s marker genes. (**c**) Bar graph showing the proportional representation of each keratinocyte sub-cluster in skin and acute wounds. (**d**) Milo analysis of keratinocyte abundance difference between Wound1 and Skin. The left panel shows the graph representation of neighborhoods. The node size and edges are proportional to the number of cells and the overlapped cell numbers between any two nodes, respectively. Nodes are colored by log2(fold changes) of cell abundance between conditions. The right panel shows a beeswarm plot representing the distribution of neighborhood abundance in cell types. Blue and red dots indicate significantly (SpatialFDR < 0.1) decreased (logFC < 0) and increased (logFC > 0) cell abundance, respectively. Color intensity indicates the degree of significance for each neighborhood. (**e**) Cellular proportions of Bas-/Spi-mig in the deconvolution of public bulk RNA-seq data (GSE174661). (**f**) Gene ontology (GO) analysis showing biological process (BP) terms enriched in basal and spinous migrating clusters. The top 200 markers (ranked by fold changes) of each cluster were used. (**g**) Dot plots showing the expression of genes associated with GO terms in (**f**). (**h**) Deconvolution of Bas-prolif, Bas-mig, and Spi-mig in acute wound ST-seq data showing distinctive migration and proliferation zones at wound edges. (**i**) Fluorescence in situ hybridization (FISH) images showing the expression of MMP3 (upper panel, Bas_mig) and KRT6B (lower panel, Spi_mig) during wound healing. Scale bar = 500um, inset plot scale bar = 200um. Significance was assessed using generalized linear modeling on a quasi-binomial distribution (**c**) and Mann-Whitney U test (**e**), *: *p < 0.05*, **: *p < 0.01*, ***: *p < 0.001*.

Upon examining the migrating keratinocytes, we found that both Bas-mig and Spi-mig clusters expressed genes essential for epithelial cell migration (e.g., *HBEGF, ANXA3, PRSS3, S100A2, FGFBP1*) and neutrophil activation (e.g., *SERPINB1/3/4, S100A7/8/9/11*), as shown by Gene Ontology (GO) analysis (**Figure 2f**). These genes peaked during the inflammatory phase (Wound1) (**Figure 2g**, **Table S3**). In the subsequent proliferative phase (Wound7), Bas-mig keratinocytes exhibited increased expression of ECM organization and cell adhesion-related genes (e.g., laminin 5, integrins, collagen), indicative of the late stages of re-epithelialization where cell-to-cell and cell-to-matrix adhesions are re-established (**Figure 2g**). Spatial deconvolution of ST-seq with paired scRNA-seq revealed Spi-mig keratinocytes overlaying Bas-mig at the wound leading edge, surrounded by proliferative keratinocytes (Bas-Prolif) (**Figure 2h**). This cellular arrangement forms rapidly post-injury, intensifies during the proliferative phase, and recedes by the remodeling phase (Wound30) (**Figure 2h**). This pattern was further confirmed in additional human wound tissues through RNA-fluorescence *in situ* hybridization (FISH) and spatial gene expression analysis (**Figure 2i**, **Figure S3d**).

To identify keratinocytes directly involved in wound repair, we sorted all scRNA-seq-analyzed keratinocytes from human skin and acute wounds into two groups using a Gaussian mixture model: 10,064 wound-associated (*KRT6A^+^*) and 17,041 non-wound (*KRT6A^-/dim^*) cells (**Figure S3e**). We targeted KRT6A expression as a marker for post-injury keratinocytes^27,28^, which was validated by pronounced KRT6A expression in wound-edge keratinocytes observed in our ST-seq analysis, in contrast to its absence in intact skin (**Figure S3f**). Pseudotemporal analysis traced a differentiation pathway from basal to spinous and then to granular keratinocytes for both wound and non-wound cells (**Figure 3a, b**). Notably, within the wound-associated group, two additional branches emerged: one transitioning from Bas-I to Bas-mig and another from Spi-II to Spi-mig, suggesting that these migrating cells originate from wound-edge keratinocytes with respective differentiation states (**Figure 3b**).

**Figure 3.**
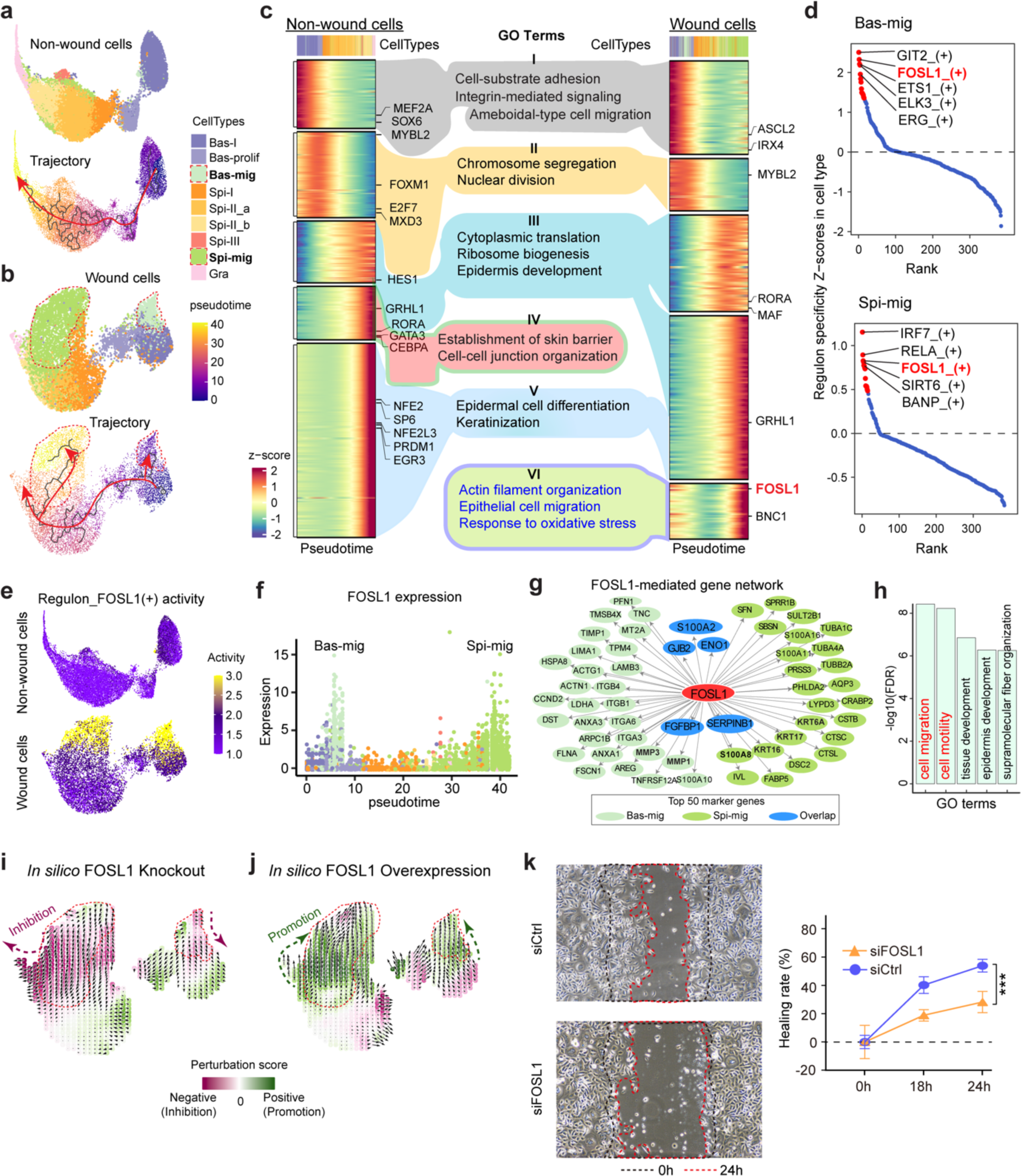
Gene network interference identifies FOSL1 as a key driver of keratinocyte migration. UMAP of cell clusters (upper panel) and pseudotime trajectories (lower panel) in non-wound (**a**) and wound keratinocytes (**b**). (**c**) Heatmaps showing expression patterns of driving genes along the pseudotime. Transcription factors (TFs) within driving genes are labeled on the right. GO terms associated with different patterns are shown in the middle. (**d**) Scatter plots showing the top 5 TF regulons based on specificity Z-scores in Bas_mig (top) and Spi_mig (bottom). (**e**) UMAP of *FOSL1* regulon activity in non-wound and wound cells. (**f**) The *FOSL1* gene expression with pseudotime. (**g**) The top 50 marker genes of Bas-mig (light green), Spi-mig (green), and both cell clusters (blue) regulated by FOSL1. (**h**) GO BP terms enriched in the FOSL1-regulated genes shown in (**g).** Perturbation simulation vector fields in wound keratinocytes with *in silico FOSL1* knockout (**i**) or overexpression (**j**). The positive (green) and negative (purple) perturbation scores indicate promotion and inhibition of cell state change, respectively. (**k**) Scratch wound assay of human primary keratinocytes with *FOSL1* expression silencing. one-way ANOVA test, \*\*\**P < 0.001*.

In summary, our study suggests a novel model of re-epithelialization in human wounds, characterized by organized keratinocyte proliferation, differentiation, and migration. Mirroring recent findings in murine skin^29,30^, we identified two distinct zones of epidermal cells around the wound: a non-proliferative migrating front surrounded by a highly proliferative hub. In contrast to murine wounds, where migrating keratinocytes often proliferate, creating a mixed zone^30,31^, human wounds show a distinct separation between keratinocyte proliferation and migration (**Figure S3g**).

### Gene network interference identifies FOSL1 as a key driver of keratinocyte migration

To compare the gene regulatory networks in various keratinocyte states during homeostasis and wound repair, we analyzed genes showing significant expression changes along pseudotemporal trajectories of wound and non-wound keratinocytes, categorizing them by their expression pattern (**Figure 3c**). GO analysis revealed that both wound and non-wound keratinocytes share several GO term patterns, including cell-substrate adhesion (I), nuclear division (II), ribosome biogenesis/epidermal development (III), and keratinization (V), indicating a common differentiation path. Notably, non-wound keratinocytes, especially towards the end of their trajectory (e.g., Spi-II/Gra), expressed genes linked to skin barrier formation and cell-to-cell junctions (IV), which were absent in wound keratinocytes. Conversely, wound keratinocytes displayed a unique expression pattern (VI) involving epithelial migration, actin organization, and oxidative stress response, prominently seen in Bas-mig and Spi-mig keratinocytes at the trajectory’s start and end, respectively (**Figure 3c**).

Combining our pseudotime analysis with SCENIC analysis^32^ to explore the gene regulatory network driving the migrating keratinocyte phenotype, we identified FOSL1 as a critical master regulator in both Bas-mig and Spi-mig clusters (**Figure 3c, d**, **Table S3**). FOSL1, a vital component of the AP1 transcriptional complex, is involved in cell differentiation, stress response, and cancer metastasis^33,34^; however, its specific role in keratinocyte behavior during wound healing was previously unclear^35^. We observed that FOSL1 exhibited high regulatory and expression specificity in these migrating cells (**Figure 3e, f**), and GO analysis of FOSL1-targeted genes in these clusters underscored their crucial role in cell migration (**Figure 3g, h**, **Table S3**).

To better understand the impact of FOSL1 perturbations on keratinocyte cell states in human wounds, we used the CellOracle package^36^. Our *in silico* analysis showed that disrupting FOSL1 impeded the migratory state of keratinocytes while enhancing FOSL1 promoted their migratory phenotype (**Figure 3i, j**). These computational predictions were experimentally validated: silencing FOSL1 with siRNA significantly reduced the motility of human keratinocyte progenitors in a scratch assay (**Figure 3k, Figure S3h**). Together, both *in silico* and experimental findings highlight FOSL1’s crucial role in regulating keratinocyte mobility.

Consistent with its role in cell migration, FOSL1 was transiently upregulated in basal and suprabasal keratinocytes at wound edges but not in intact skin, as shown by our scRNA-seq and ST-seq data, and confirmed by immunofluorescence (IF) staining (**Figure 6b, c, Figure S3i**). This upregulation also occurred in migrating keratinocytes in a mouse acute wound model, suggesting an evolutionarily conserved role in keratinocyte migration (**Figure 6d**, **Figure 7b, c**). Notably, this increase in FOSL1 expression was absent in other inflamed skin conditions, such as psoriasis and atopic dermatitis, underscoring its specific involvement in wound repair (**Figure S3j**)^16,37^.

### Pro-inflammatory macrophages support re-epithelialization at the inflammatory phase

Our combined scRNA-seq and ST-seq analysis extends beyond keratinocytes, revealing the dynamic gene expression and cellular diversity of immune cells, fibroblasts, and angiogenic cells throughout human skin wound healing. We identified 11 myeloid cell types in human skin and acute wounds, including four macrophage clusters: Mac_inf (*APOE^+^*, *CXCL1^+^)*, Mac1 (*IL1B*^+^, *THBS1^+^*, *EREG*^+^), Mac2 (*DAB2^+^*, *C1QA/B^+^)*, and Mac3 (*MMP19^+^*, *MMP9^+^*, *VEGFA^+^*)^38,39^; four dendritic cell clusters: plasmacytoid DC (pDC, *ACOT7^+^*, *LTB^+^*, *IGKC^+^*), conventional DC1 (cDC1, *CLEC9A^+^*, *WDFY4*), cDC2 (*CD1C^+^*, *IL1R2^+^*, *CLEC10A^+^*), and DC3 (*CCR7^+^, LAMP3^+^*); Langerhans cells (LC: *CD207^+^*, *CD1A^+^*); and subsets of apoptotic (*DNAJB1^+^, HSPA1B^+^*) and cycling (*PCLAF^+^, H4C3^+^*) cells (**Figure 4a, b**, **Figure S4a**, **Table S4**). Mac_inf and Mac1 are more pro-inflammatory, while Mac2 and Mac3 are more anti-inflammatory, reflecting M1 to M2 transition signatures^40^ (**Figure S4b**). Additionally, using a modified Cell Ranger pipeline, we identified neutrophils *(CSF3R^+^*, *FCGR3B^+^*, *CXCR2^+^*, and *CMTM2^+^)*^41^, which were initially missed in scRNA-seq analysis due to their low RNA content and high RNase levels (**Figure S4c-g**). We also identified nine clusters of lymphoid cells, including regulatory T cells (Treg: *TIGIT^+^*, *BATF^+^*, *FOXP3^+^*), helper T cells (Th: *LDHB^+^*, *KLF2^+^*, *GIMAP7^+^*), innate lymphoid cells (ILC: *AHR^+^*, *CCR6^+^*, *PTGER4^+^*), cytotoxic T cells (Tc: *TRGC2^+^*, *KLRC2/3^+^*), ILC1/natural killer cells (*XCL1/2^+^, FCER1G^+^*), NK cells (*GZMA/K^+^*), tolerant T cells (Ttol: *DNAJB1^+^*, *NR4A1^+^*)^42^, plasma cells (*PTGDS^+^*, *JCHAIN^+^*, *IL3RA^+^*), and B cells (*IGHM^+^*, *MS4A1^+^*, *CD79A^+^*)^16,42,43^ (**Figure 4f, g**, **Figure S4h**, **Table S4**).

**Figure 4.**
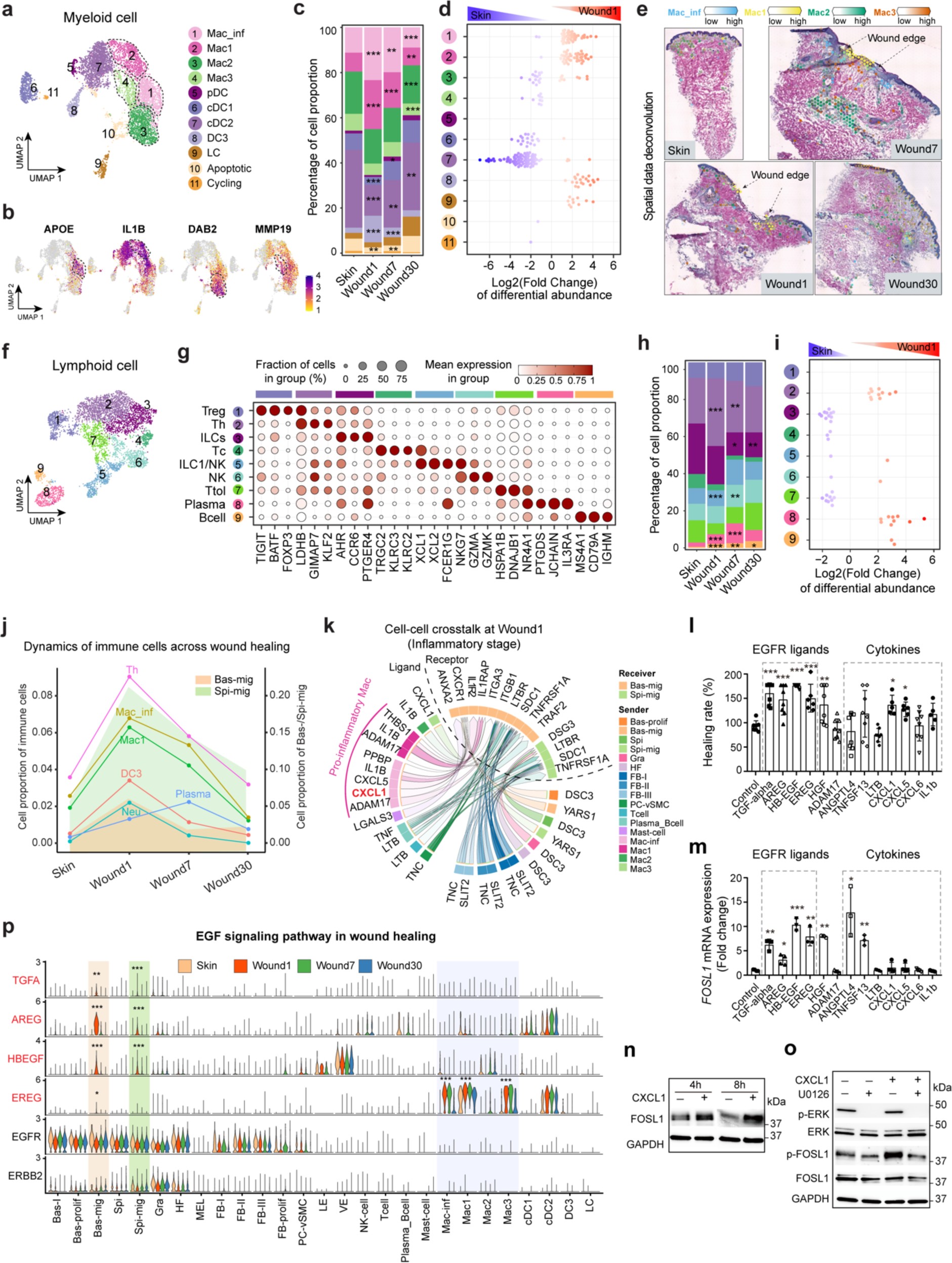
Pro-inflammatory macrophages support re-epithelialization at the inflammatory phase. (**a**) UMAP of myeloid subpopulations in human acute wounds. Mac_inf, inflammatory macrophage; pDC, plasmacytoid dendritic cell; cDC1/2, conventional DC1/2; LC, Langerhans cell. (**b**) Feature plots showing macrophage markers of APOE (Mac_inf), IL1B (Mac1), DAB2 (Mac2), and MMP19 (Mac3). (**c**) Bar graph showing the proportional representation of myeloid cell types during wound healing. (**d**) Milo beeswarm plot showing the differential abundance of cell types between Wound1 and Skin. Blue and red dots indicate significantly (SpatialFDR < 0.1) decreased (logFC < 0) and increased (logFC > 0) cell abundance, respectively. Color intensity indicates the degree of significance for each neighborhood. (**e**) Deconvolution of macrophages in wound healing ST-seq data. (**f**) UMAP of lymphoid subpopulations in acute wounds. (**g**) Dot plot showing marker gene expression of each cell cluster. Treg, regulatory T cell; Th, helper T cell; ILCs, innate lymphoid T cell; Tc, cytotoxic T cell; NK, natural killer T cell; Ttol, tolerant T cell. (**h**) Bar graph showing the proportional representation of lymphoid cell types. (**i**) Milo beeswarm plot showing the differential abundance of cell types between Wound1 and Skin. (**j**) Summary plot illustrating the dynamics of immune cells and migrating keratinocytes during wound healing. (**k**) Circos plot showing the top 50 cell-cell interactions between all cell types (Ligand) and migrating keratinocytes (Receptor) in Wound1. Cell migration assay (**l**) and *FOSL1* mRNA expression (**m**) in primary human keratinocytes treated with growth factors or cytokines. (**n**) Western blot of FOSL1 protein in keratinocytes treated with CXCL1 for 4 and 8 hours. (**o**) Western blot of phosphorylated and total ERK and FOSL1 proteins in keratinocytes treated with CXCL1 for 30 minutes, with or without ERK pathway inhibitor U0126. GAPDH was used as a loading control. (**p**) Violin plot showing EGF signaling ligand and receptor expression in each cell type during skin wound healing. Significances were determined using generalized linear modeling on a quasi-binomial distribution (**c**, **h**), One-way ANOVA test (**l**, **m**), and Mann-Whitney U test (**p**), comparing other conditions with normal skin/control group, *: *p < 0.05*, **: *p < 0.01*, ***: *p < 0.001*.

Cell proportion and Milo^25^ analyses highlighted several immune cells peaking during the inflammatory phase of wound repair: (1) pro-inflammatory macrophages (Mac_inf and Mac1) located in the upper dermis adjacent to migrating epithelial cells (**Figure 4c-e**); (2) neutrophils, known as the first myeloid cells recruited from circulation (**Figure S4i, j**); (3) DC3 cells, noted for their maturity and migratory capabilities, with significant cytokine^44^ (**Figure 4c, d**); and (4) Th cells, characterized by high expression of KLF2, a transcription factor that regulates T-cell migration and differentiation^45^ (**Figure 4h, i**, **Figure S4k**). Plasma and B cells also increase during the inflammatory phase and peak during the proliferative phase (**Figure 4h, i**). Conversely, cDC1 and cDC2 proportions initially decrease at the onset of inflammation but rebound as healing progresses, paralleling the reduction in neutrophils and macrophages (**Figure 4c, d**).

During the inflammatory phase of wound repair, keratinocytes began migrating rapidly, coinciding with the peak activity of immune cells (**Figure 4j**). To explore whether immune cells signal keratinocytes to promote re-epithelialization, we used the MultiNicheNet R package^46^ to analyze cell-to-cell communication. We focused on the top 50 ligand-receptor interactions influencing migrating keratinocytes, which included many signals previously known to enhance keratinocyte motility, such as THBS1^47^, LGALS3^48^, and TNF^49^ (**Figure 4k**, **Figure S5a**, **Table S5**). Notably, CXCL1 and CXCL5, typically involved in recruiting inflammatory cells^50^, also promoted keratinocyte migration in a FOSL1-dependent manner (**Figure 4l**, **Figure S5b, c**). While CXCL1 did not alter FOSL1 mRNA levels (**Figure 4m**), it triggered the phosphorylation of Ser265 in FOSL1’s C-terminal destabilizer region, enhancing FOSL1 stability^51^ (**Figure 4n, o**, **Figure S5d**). This effect was reversed by the ERK pathway inhibitor U0126, highlighting the role of ERK signaling in this regulatory process (**Figure 4o**).

Furthermore, using the CellChat package^52^, we compared cell-to-cell communication between skin and wounds, identifying EGFR signaling as a wound-specific pathway influencing both Bas-mig and Spi-mig keratinocytes (**Figure 4p**, **Figure S5e, f**). Intriguingly, while EGF receptors (*EGFR* and *ERBB2*) were highly expressed on keratinocytes and fibroblasts, the expected EGF ligand was absent in our scRNA-seq and ST-seq data, questioning its assumed role in wound healing^53^. Instead, ligands such as TGFA, AREG, and HB-EGF were rapidly upregulated in migrating keratinocytes, likely serving as autocrine signals during the inflammatory phase, whereas EREG was predominantly expressed by wound macrophages (**Figure 4p**). These EGFR ligands significantly enhanced keratinocyte migration and induced mRNA expression of FOSL1, a key regulator of keratinocyte motility (**Figure 4l, m**, **Figure S5b**).

Considering the cellular sources of these pro-migratory signals, we analyzed the spatial co-occurrence of cell types in our ST-seq data, noting that cell-to-cell communication, particularly juxtacrine and paracrine signaling, is spatially restricted (**Figure 1f**). In niches with migrating keratinocytes (niche7 and niche8), we observed their close associations with pro-inflammatory macrophages and plasma cells at the wound edge (**Figure 1f**, **Figure S2e**), further confirmed by IF staining (**Figure 6c**).

Our study thus outlines a detailed framework for understanding the cell-to-cell communication signals directing re-epithelialization: in the early inflammatory phase, keratinocytes generate autocrine EGF signals (TGFA, AREG, HBEGF) while nearby pro-inflammatory macrophages contribute paracrine EGF (EREG) and chemokine signals (CXCL1 and CXCL5), collaboratively enhancing FOSL1 expression at both mRNA and protein levels and promoting keratinocyte migration.

### Fibroblasts play a major role in promoting re-epithelialization at the proliferative phase

Our scRNA-seq identified four main fibroblast (FB) clusters consistent with previous studies^9,54^: mesenchymal (FB-I: *POSTN^+^*), pro-inflammatory (FB-II: *C3^+^*), papillary (FB-III: *ELN^+^LEPR^+^*), and proliferating (FB-prolif: *MKI67^+^*) FBs (**Figure 5a-c**, **Figure S6a**, **Table S4**). FB-I was subdivided into four subclusters with distinct markers (*COL11A1^+^*, *MMP11^+^*, *COL4A1^+^*, and *SFRP4^+^COMP*^+^), while FB-II was split into two subclusters differentiated by apolipoproteins (*APOD^+^*or *APOE^+^*) and immune genes (*ITM2A^+^* or *CCL19^+^*). Additionally, two new FB clusters were identified: one adjacent to hair follicles (*SFRP1^+^CRABP1^+^*) and another similar to papillary FBs (*ELN^+^SFRP4^+^*), as shown in ST-seq deconvolution and dendrogram analysis, respectively (**Figure 5c, Figure S6b**). RNA velocity analysis^55^ depicted two FB trajectories: one from FB-I(*SFRP4^+^COMP^+^*) to FB-II(*APOE^+^CCL19^+^*) and another from FB-I(*POSTN^+^COL11A1^+^*) to FB-III(*ELN^+^LEPR^+^*) (**Figure S6c**). Cells in the initial state of the first trajectory highly expressed PI16, a progenitor fibroblast marker^56^ (**Figure S6d**). Cellular proportion and Milo analysis showed that proliferating fibroblasts were specifically present during the proliferative phase (Wound7), mesenchymal fibroblasts (FB-I) increased and dominated the wound bed in the remodeling phase (Wound30), and pro-inflammatory (FB-II) and papillary (FB-III) fibroblasts declined as healing progressed (**Figure 5d, e, Figure S6e**). These shifts in fibroblast heterogeneity were validated by ST-seq deconvolution and further confirmed by FISH (**Figure 5f, g**).

**Figure 5.**
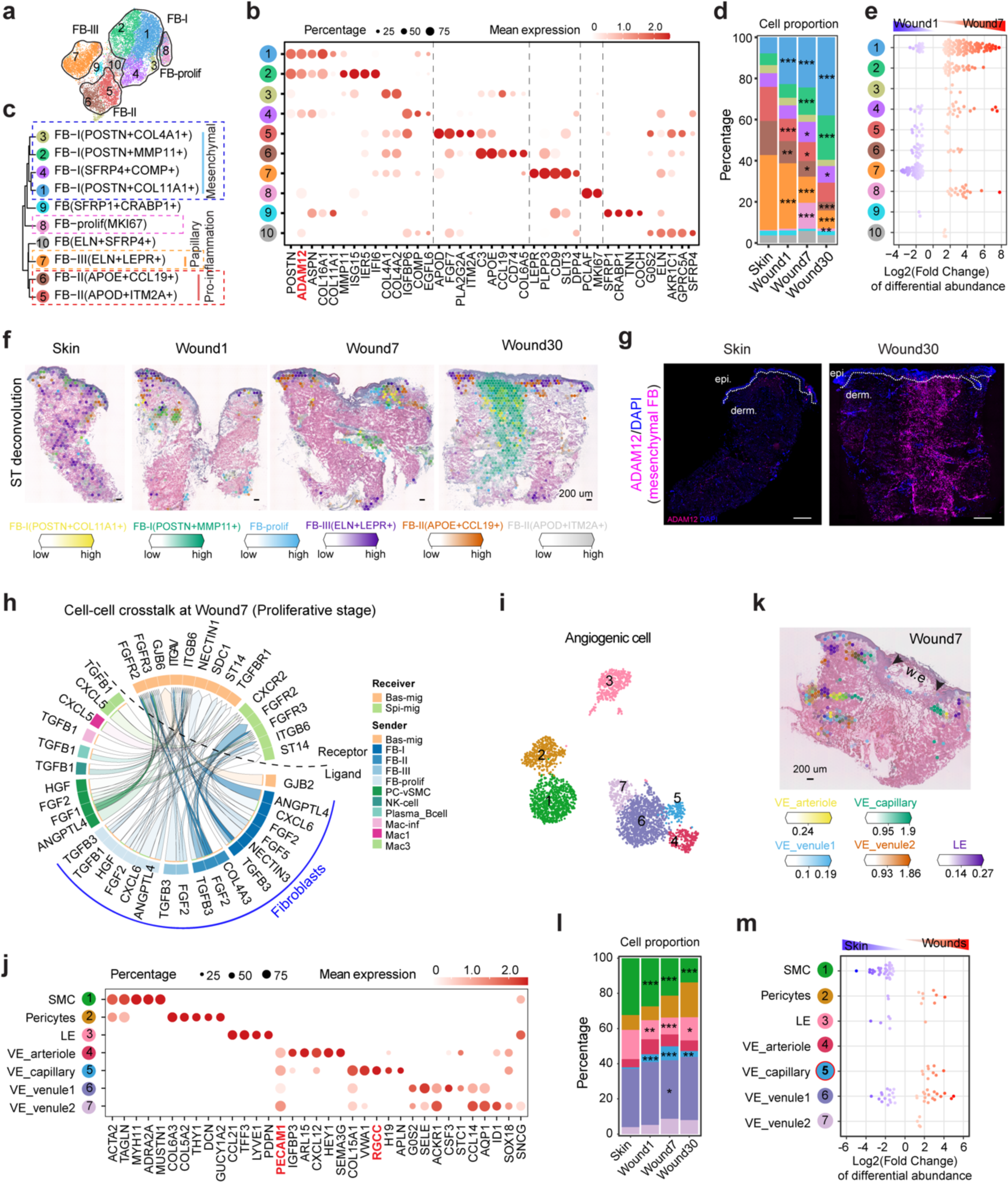
Fibroblasts play a major role in promoting re-epithelialization at the proliferative phase. (**a**) UMAP of fibroblast subpopulations in acute wounds. (**b**) Dot plot showing marker gene expression of FB clusters. (**c**) The dendrogram illustrates cell clusters’ relatedness based on a distance matrix constructed in gene expression space. FB-, fibroblast; FB-prolif, proliferating fibroblasts. (**d**) Bar graph showing the proportional representation of FB cell types during wound healing. (**e**) Milo beeswarm plot showing the differential abundance of cell types between Wound7 and Wound1. Blue and red dots indicate significantly (SpatialFDR < 0.1) decreased (logFC < 0) and increased (logFC > 0) cell abundance, respectively. Color intensity indicates the degree of significance for each neighborhood. (**f**) Deconvolution of fibroblast subpopulations in wound healing ST-seq data. (**g**) Fluorescence in situ hybridization (FISH) images of ADAM12 expression (a marker for mesenchymal fibroblast: FB-I) in Skin and Wound30. Scale bar = 500um. (**h**) Circos plot showing the top 50 cell-cell interactions between all cell types (Ligand) and migrating keratinocytes (Receptor) in Wound7. (**i**) UMAP of angiogenic subpopulations in acute wounds. (**j**) Dot plot showing marker gene expression of cell clusters. SMC, smooth muscle cell; LE, lymphatic endothelium; VE-, vascular endothelium. (**k**) Deconvolution of endothelial populations in Wound7 ST-seq data. (**l**) Bar graph showing the proportional representation of angiogenic cell types during wound healing. (**m**) Milo beeswarm plot showing the differential abundance of angiogenic cell types between Wounds and Skin. Significance was assessed using generalized linear modeling on a quasi-binomial distribution (**d**, **l**), comparing other conditions to normal skin, *: *p < 0.05*, **: *p < 0.01*, ***: *p < 0.001*.

Upon comparing the signals influencing keratinocyte migration across the inflammatory and proliferative phases, we noted a shift from macrophages as primary influencers during the inflammatory phase to fibroblasts playing a pivotal role in the proliferative phase (**Figure 4k, 5h**, **Figure S6f**). Proliferating fibroblasts increased the production of growth factors such as HGF, FGF2, and TGFβ1 during the proliferative phase, boosting keratinocyte motility^57–59^ (**Figure 5h**, **Figure S6f**, **Figure 4l**, **Table S5**). HGF also upregulated FOSL1 expression in keratinocytes, a critical factor in cell motility (**Figure 4m**, **Table S5**). Additionally, ST-seq data revealed close associations between proliferating fibroblasts and Bas-mig keratinocytes (niche7), as well as inflammatory macrophages (niche13) at the wound edge, highlighting fibroblasts’ role in re-epithelialization (**Figure 1f**, **Figure S2e**). Thus, our findings suggest that pro-inflammatory macrophages and fibroblasts sequentially support keratinocyte migration during different healing stages, functioning like a relay race.

Our scRNA-seq analysis also identified seven angiogenic cell types in human skin and wounds, including various endothelial cells (lymphatic, arteriole, and capillary) and two venule endothelial subsets, along with associated smooth muscle cells and pericytes (**Figure 5i, j**, **Table S4**). Deconvolution of ST-seq data highlighted well-defined vascular structures formed by these cells in the dermis (**Figure 5k**, **Figure S6g, h**). Post-injury, cellular proportion and Milo analysis showed a decrease in smooth muscle cells and lymphatic endothelial populations, while capillary endothelial cells notably increased during the proliferative phase, indicating active angiogenesis (**Figure 5l, m**). Additionally, ST-seq data revealed close associations between capillary endothelial cells, proliferating fibroblasts, and pro-inflammatory macrophages (niche13) during the proliferative phase, characterizing the newly formed granulation tissue (**Figure 1f**, **Figure S2e**).

### Multi-facet pathological changes in chronic wounds

To advance our research on chronic wound pathology, we integrated scRNA-seq data from acute human wounds with VUs and DFU data^8^ using Harmony^24^ (**Figure 1g**). We identified distinct pathological changes between chronic wound types, differing by prognosis (DFU_Healing vs. DFU_NonHealing) and etiology (VU vs. DFU).

We found significant reductions in migratory keratinocytes in DFUs and their complete absence in VUs, correlating with re-epithelialization failure in chronic wounds^3,60^ (**Figure 6a**). Consistent with this, scRNA-seq data revealed fewer FOSL1^+^ keratinocytes in both DFUs and VUs compared to acute wounds, a trend more evident at the protein level as shown by IF analysis (**Figure 6b, c**). In a mouse model, increased FOSL1 expression was observed at the wound edges of normal mice but not in diabetic (db/db) mice, indicating that FOSL1 deficiency hinders re-epithelialization (**Figure 6d**). Unlike acute wounds, where keratinocyte proliferation increases (**Figure S3c**, **Figure S7a**), non-healing DFUs show reduced keratinocyte proliferation, while VUs display highly proliferative keratinocytes at wound edges, consistent with the hyperproliferative epidermis observed in VU edges^61^ (**Figure S7a**).

**Figure 6.**
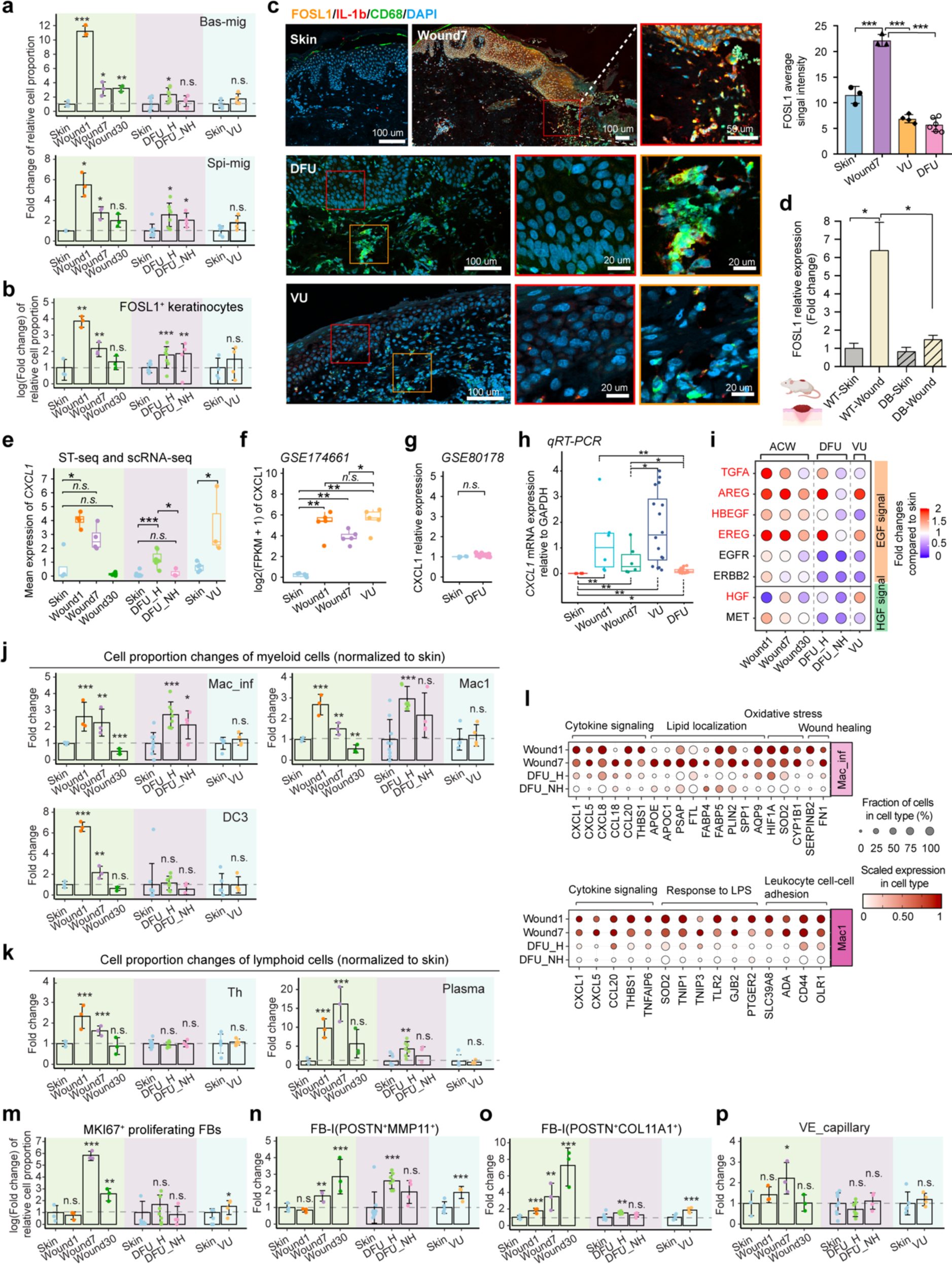
Multi-facet pathological changes in chronic wounds. (**a**) Bar charts showing fold changes of cell proportions of Bas-mig (upper panel) and Spi-mig (lower panel) in acute and chronic wounds by normalized to healthy skin. Bas-/Spi-mig, basal/spinous migrating keratinocytes, DFU_H/_NH, healed/non-healed diabetic foot ulcers, VU, venous ulcer. (**b**) Bar charts showing the log2(fold change) of FOSL1^+^ cell proportion in total keratinocytes normalized to skin. (**c**) Immunofluorescence staining of FOSL1 (migrating keratinocyte marker), IL1b, and CD68 (pro-inflammatory macrophage markers) in human skin, acute, and chronic wound edges. DAPI stains the nuclei. Scale bars: 100um in low-magnification images, 20um in high-magnification images. The signal intensity of epidermal FOSL1 was quantified. (**d**) Relative gene expression of FOSL1 in the epidermis of wild type (WT) and diabetic (DB) mouse skin and wounds. n= 5. Box plots showing CXCL1 expression in our ST-seq (acute wounds) and scRNA-seq (DFU and VU) datasets (**e**), public bulk RNA-seq data of skin and acute wounds from 5 donors and 5 VUs (**f**), and microarray data of skin (n=6) and DFU (n=6) (**g**). (**h**) qRT-PCR of CXCL1 in human skin and acute wounds from 7 donors, 16 VUs, and 27 DFUs. (**i**) Dot plot showing the fold changes of ligands and receptors of EGF and HGF signals in acute (ACW) and chronic wounds (DFU and VU) normalized to the control skin. Bar charts showing fold changes of cell proportions of myeloid cells (**j**), lymphoid cells (**k**), proliferating FB (**m**), FB-I(POSTN^+^MMP11^+^) (**n**), FB-I(POSTN^+^COL11A1^+^) (**o**), and VE-capillary (**p**) in acute and chronic wounds normalized to control skin. Mac_inf, inflammatory macrophage; DC, dendritic cell; Th, T helper cell; FB, fibroblast; VE, vascular endothelial cell. (**l**) Dot plots showing scaled expression of differentially expressed genes between acute wounds and DFU in Mac_inf (upper panel) and Mac1 (lower panel). Genes enriched in relevant GO terms were plotted. Significances were assessed using generalized linear modeling on a quasi-binomial distribution (**a**, **b**, **j**, **k**, **m-p**), One-way ANOVA test (**c**, **d**), and Mann-Whitney U test (**e-h**), *: *p < 0.05*, **: *p < 0.01*, ***: *p < 0.001*, n.s.: no significance.

In human chronic wounds, we also analyzed cell-to-cell communication signals crucial for re-epithelialization, including CXCL1, EGFR ligands, and HGF. Analysis using single-cell data (*Theocharidis, G*. *et al.*^8^ and ours), bulk RNA-seq (GSE174661^26^), and microarray data (GSE80178^62^) showed that CXCL1 was upregulated in VUs, similar to acute wounds, but not in DFUs (**Figure 6e-g**). qRT-PCR confirmed these results, indicating higher CXCL1 levels in VUs and lower in DFUs (**Figure 6h**). Additionally, scRNA-seq data demonstrated abundant EGF ligands and receptors expression in acute wounds, but this was reduced or weakly induced in non-healing DFU and VU compared to normal skin (**Figure 6i**). HGF and its receptor MET also displayed low expression in DFUs, while in VUs, their levels were similar to those in acute wounds (**Figure 6i**). Therefore, lacking FOSL1^+^ migrating keratinocytes in VUs may be linked to inadequate EGF signaling, while diminished CXCL1, EGF, and HGF signals may collectively hinder healing in DFUs.

Our research highlights the critical role of inflammation in tissue repair^63^. In contrast to the significant increase of pro-inflammatory Mac, DC3, plasma cells, and Th cells in acute wounds, these cells were notably scarce in DFUs and VUs. Specifically, VUs exhibited a marked deficiency in pro-inflammatory macrophages (**Figure 6j, k**, **Figure S7b**). Although DFUs maintained similar proportions of Mac_inf and Mac1 as acute wounds, their macrophages displayed a significant reduction in gene expression crucial for cellular functions^64^, such as cytokine signaling (CXCL1, CXCL5, CCL20), lipopolysaccharide (LPS) response, and oxidative stress (**Figure 6l**). These findings suggest an impaired inflammatory response in chronic wounds, which may revise the conventional view of sustained inflammation in these conditions^65,66^.

Furthermore, we observed a notable absence of proliferating and mesenchymal fibroblasts in chronic wounds, suggesting a deficiency in fibroblast proliferation and a reduced mesenchymal stem cell response^67,68^ (**Figure 6m-o**). Neither DFUs nor VUs exhibited the expected increase in VE_capillary cells characteristic of the proliferative phase of acute wounds, suggesting impeded angiogenesis, in line with previous studies^69^ (**Figure 6p**). Specifically, VUs displayed increased venule endothelial cells, reflecting the vascular abnormalities associated with chronic venous insufficiency^70^ (**Figure S7c**).

By comparing scRNA-seq data from human acute wounds with chronic wounds, we identified critical pathological changes, including compromised re-epithelialization, altered inflammatory responses, impaired granulation tissue formation, and hindered angiogenesis. Notably, DFUs with better healing outcomes displayed less severe pathological changes, characterized by increased presence of Bas-mig keratinocytes, inflammatory macrophages, mesenchymal fibroblasts, and enhanced CXCL1, EGF, and HGF signaling, highlighting the critical role of these processes in wound healing. Our data thus provide a valuable human *in vivo* gene expression framework to further investigate the molecular mechanisms driving these pathological changes and to develop targeted therapeutic strategies.

### Comparison of human and murine skin wound healing

While studying cellular heterogeneity in skin wounds of mouse models has enhanced our understanding of wound healing dynamics^19,71–74^, human wound healing mechanisms differ significantly due to variations in skin structure and healing processes^75^. To bridge this gap, we directly compared human and mouse wounds by integrating scRNA-seq data from both species using canonical correlation analysis (CCA) (**Figure 7a, Figure S8a**)^14^. We identified similar cell types across species, such as migrating keratinocytes with shared markers like NRG1, IL24, FOSL1, AREG, and GJB2, suggesting a conserved regulatory mechanism for keratinocyte migration (**Figure 7a-c**, **Table S6**). Notably, recent research has shown that IL-24, upregulated in epithelial stem cells, promotes wound repair by enhancing re-epithelialization, vascular regeneration, and fibroblast activation, acting independently of microbial and adaptive immune factors^76^.

**Figure 7.**
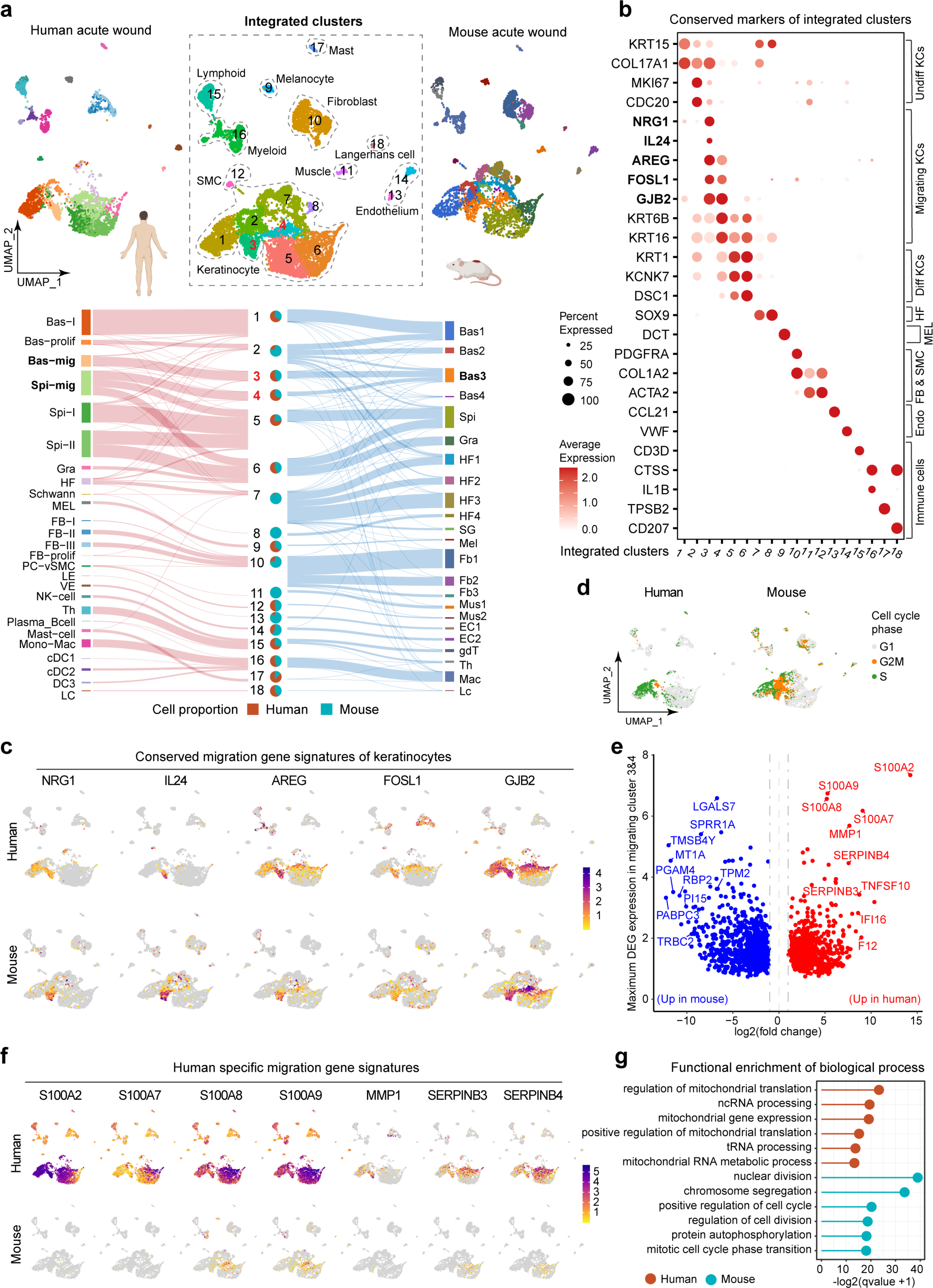
Comparison of human and murine skin wound healing. (**a**) UMAP of integrated cell clusters from human and mouse acute wounds. The top-left and top-right plots show color-coded human and mouse cell types separately. The Sankey diagram traces the cellular assignments of integrated clusters from the original human and mouse cell types, and the pie chart reveals their contributions to each cluster. Bas-/Spi-/Gra-, basal/spinous/granular keratinocyte (KC); Bas-prolif, proliferating keratinocyte; Bas-/Spi-mig, migrating keratinocyte; HF, hair follicle; MEL/Mel, melanocyte; FB/Fb, fibroblast; PC-vSMC, pericyte and vascular smooth muscle cell; LE, lymphatic endothelium; VE, vascular endothelium; NK-cell, natural killer cell; Th, T helper cell; Mono-mac, monocyte and macrophage; cDC, conventional dendritic cell; LC, Langerhans cell; SG, sweat gland cell; Mus, muscle cell; EC, endothelial cell; gdT, gamma-delta T cell. (**b**) Dot plot showing the expression of conserved markers of integrated clusters. Undiff/Diff, undifferentiated/differentiated KC; Endo, endothelium. (**c**) Feature plots showing representative conserved gene signatures of migrating keratinocytes. (**d**) Cell cycle analysis in human and mouse wound cells. (**e**) Volcano plot showing differentially expressed genes (DEGs) in migrating keratinocytes (clusters 3&4) between human and mouse wounds. Red and blue dots represent upregulated genes in humans and mice, respectively. The top 10 changed genes are highlighted. (**f**) Feature plots showing representative human-specific migration gene signatures. (**g**) Lollipop chart showing enriched GO terms of human- and mouse-DEGs in (**e)**.

Our scRNA-seq data reveal distinct structural differences between murine and human skin: humans have a thicker epidermis with more spinous keratinocytes (integrated clusters 4-6), mice possess abundant hair follicles (clusters 7 and 8) and a unique panniculus carnosus structure (cluster 11 MYF6/Mrf4^+^)^77^ (**Figure 7a, Figure S8b**). These structural variances affect wound healing mechanisms: mice primarily heal through panniculus carnosus contraction, whereas humans rely on re-epithelialization and granulation tissue formation^78^. Additionally, murine wounds show a higher presence of fibroblasts (cluster 10) and endothelial cells (clusters 13 and 14), while human wounds exhibit more mast cells (cluster 17), aligning with findings of greater mast cell heterogeneity in humans than in mice^79^. Murine wounds also have more proliferating cells at the G2/M phase of the cell cycle, consistent with their more robust healing capabilities (**Figure 7d**).

Beyond cellular composition, gene expression differences between mice and humans were even more striking (**Figure 7e**, **Table S6**). For instance, migration-related genes such as MMP1, S100A2/7/8/9, and SERPINB3/4 are highly expressed in humans but barely in mice (**Figure 7e, f**). MMP1 supports human keratinocyte migration by breaking down dermal collagen, a role filled by MMP13 in mice, indicating species-specific proteolysis mechanisms for re-epithelialization^80,81^. S100A proteins contribute to antimicrobial defense and tissue repair^82^, while SERPINB3/4 play a crucial role in keratinocyte inflammatory responses^83^. The distinct gene expressions likely reflect differing adaptations to microbial threats. Further GO analysis of DEGs showed that human migrating keratinocytes are enriched with genes related to mitochondrial activity, which support wound healing functions such as energy provision and inflammation regulation^84^ (**Figure 7g**, **Table S6**). In contrast, murine migrating keratinocytes exhibit a proliferation-centric gene expression profile, reflecting their combined roles in both cell proliferation and migration^30^ (**Figure 7g**, **Figure S3g**). Additionally, cell-to-cell communication during wound healing also varies between species; for example, EGFR ligands in human wounds are produced by macrophages, dendritic cells, and endothelial cells in addition to migrating keratinocytes (**Figure 4p**), whereas in mouse wounds, these ligands are predominantly expressed in keratinocytes and fibroblasts, but not immune cells^85^. These findings underline the limitations of using mouse models to study the human wound healing process.

Furthermore, our scRNA-seq dataset provides a unique platform to investigate human-specific genes in wound repair, identifying two protein-coding genes (IL32 and ARHGEF35)^86^ and 49 non-conserved human long non-coding RNAs^87^ with cell-type specific expression in human wounds (**Figure S8c**). Although most of these lncRNAs have demonstrated functionality in other tissues and diseases, their roles in skin and wound healing remain to be investigated. Advancing research in this area could reveal human-specific repair mechanisms pertinent to wound pathology.

In summary, our study shows that although humans and mice share many wound healing processes, there are significant differences in cellular diversity and gene expression. This underscores the importance of assessing the clinical relevance of mouse model data against the spatiotemporal roadmap of human skin wound healing our study provides.

## DISCUSSION

Wound healing involves intricate coordination of various cell types and molecular signals, which have been primarily studied in animal and *in vitro* models. To better understand how these mechanisms apply to humans, we utilized single-cell and spatial transcriptomics in a unique *in vivo* human wound healing model. This innovative approach allows us to trace the healing process’s dynamics and cellular changes with unprecedented precision and consistency across an individual’s recovery. With this spatiotemporal cell atlas of human skin wound healing, we delve into re-epithelialization, illuminating the cellular architecture of the human wound tongue, its gene regulatory networks, and the cell-to-cell communication that promotes keratinocyte motility. The study also highlights cellular and molecular discrepancies in chronic wounds and identifies potential therapeutic targets. This pivotal dataset is a vital resource for validating the relevance of animal model findings and stimulates further research into human wound healing mechanisms. To facilitate global research collaboration and drive further discoveries, we have made this groundbreaking roadmap of human skin wound healing accessible for interactive exploration online (https://www.xulandenlab.com/tools).

Cell-to-cell communication signals are promising therapeutic targets for enhancing tissue repair. Our study on human wounds challenges established paradigms, particularly in ligand-receptor signaling. Keratinocytes are pivotal in initiating immune responses through cytokines, chemokines, and growth factors, and their activity varies with the stimulus and context^88^. We identified IL18, CCL27, and CXCL14 as the primary cytokines/chemokines produced by keratinocytes in acute human wounds (**Figure S8d**). IL-18 is critical in starting inflammation during bacterial infections^89^, CCL27 aids T cell-mediated inflammation^89^ and skin regeneration^90^, and CXCL14 recruits immune cells, inhibits angiogenesis, and has antimicrobial properties^91–93^. Despite their significant roles, these cytokines are often overlooked in wound repair studies. Additionally, some cytokines are produced by specific keratinocyte clusters (**Figure S8d**). For example, basal migrating keratinocytes express IL-20 and IL-24 targeting the IL-22R receptor; granular keratinocytes produce IL36G and IL36RN, supporting wound healing^94,95^. Furthermore, our findings reveal that, while IL1A/B/RN, IL6, and CXCL1/5/8 are commonly associated with keratinocytes *in vitro*, in human wounds *in vivo*, these cytokines and chemokines are primarily produced by macrophages, dendritic cells, and fibroblasts, not by keratinocytes (**Figure S8d**). This insight challenges previous assumptions based on *in vitro* studies and highlights the complexity of cellular interactions in actual wound environments.

Many molecular signals have been previously reported to regulate keratinocyte motility, and our study evaluated the physiological relevance of these signals in human skin wound repair. During the early inflammatory phase, keratinocytes initiate autocrine EGFR signaling, while pro-inflammatory macrophages enhance cell migration by contributing additional EGFR and CXCL1 signals. Although CXCL1 is typically associated with recruiting inflammatory cells such as neutrophils, it also plays a critical role in promoting keratinocyte migration^90^, and blocking its receptor, CXCR2, can impede re-epithelialization independently of neutrophils^96^. Our observations of reduced pro-inflammatory macrophage activity in chronic wounds align with the longstanding theory that inflammation is essential for tissue repair^63^. Actually, the current perspective on inflammation in chronic wounds is shifting from being viewed as persistently excessive^97^ to dysfunctional, characterized by impaired monocyte recruitment and macrophage and neutrophil dysfunction^98,99^. Therefore, precise modulation of pathological inflammation, rather than its inhibition, may be crucial for effective chronic wound therapy.

### Limitations of study

The Visum ST data lack single-cell resolution, and we used computational methods to deconvolve ST spots into cell-type proportions. Higher-resolution ST in future studies could shed light on the spatial arrangement of cell states at the cellular or subcellular level. Additionally, our sampling of human acute wounds at days 1, 7, and 30 post-injury captures key healing phases, but the limited time points may overlook transient cell states and ephemeral cell-to-cell cell interactions; hence, a higher temporal resolution would be advantageous. Moreover, limited tissue availability and the high costs of single-cell and spatial omics technologies constrained the number of wound samples we could study. Given the variability of human chronic wounds, more extensive cohort studies and cross-dataset comparisons are crucial to confirm our findings and aid in the molecular stratification of complex chronic wounds.

## Supporting information

STAR methods

## ACKNOWLEDGMENTS

We thank all the healthy donors and patients who participated in this study. The computations and data handling were enabled by resources in projects of sens2020010, SNIC2022/22-873, NAISS 2023/22-935 provided by the National Academic Infrastructure for Supercomputing in Sweden (NAISS) at UPPMAX, funded by the Swedish Research Council through grant agreement no. 2022-06725. This work was supported by Swedish Research Council (Vetenskapsrådet) grants (2020-01400), Ragnar Söderbergs Foundation (M31/15), Welander and Finsens Foundation (Hudfonden), Ming Wai Lau Centre for Reparative Medicine, LEO Foundation, Cancerfonden, Åke Wiberg Foundation, Karolinska Institutet, National Natural Science Fund for Excellent Young Scientists Fund of China, National Natural Science Foundation of China (82272294), the Distinguished Medical Expert of Jiangsu Province, Non-profit Central Research Institute Fund of Chinese Academy of Medical Sciences (2022-RC320-02, 2021-RC320-001, 2020-RC320-003), and CAMS Innovation Fund for Medical Sciences (2021-I2M-1-059).

## AUTHOR CONTRIBUTIONS

N.X.L., Z.L., and D.L. conceived and designed the study. P.S., L.G., J.G., C.C., Y.X., W.H., and L.Zhu. collected clinical samples. D.L., X.B., L.Zhang., L.Li., and J.W. performed experiments. Z.L. performed bioinformatics analysis with the support from Å.K.B, L.Luo., and Y.C.. M.H., M.K., C.L.B., H.G., and K.A. helped with data analysis and presentation. Z.L. and N.X.L. wrote the manuscript with input from all authors. All authors read and approved the final version of the manuscript.

