## Supplementary material for "Spatiotemporal Single-Cell Roadmap of Human Skin Wound Healing": STAR methods

### KEY RESOURCES TABLE

| REAGENT or RESOURCE | SOURCE | IDENTIFIER |
|---|---|---|
| **Antibodies** | | |
| FOSL1 | CellSignaling Technology | #5281 |
| p-FOSL1 | CellSignaling Technology | #3880 |
| ERK | CellSignaling Technology | #4695 |
| p-ERK | CellSignaling Technology | #4370 |
| GAPDH | HUABIO | ET1601-4 |
| Fra1 Polyclonal Antibody | Thermofisher | #PA5-40361 |
| Alexa Fluor® 488 Mouse monoclonal [KP1] to CD68 | Abcam | ab222914 |
| Anti-IL-1 beta antibody [OTI3E1] | Abcam | ab156791 |
| MMP3 probe | ACD company | #403421 |
| KRT6B probe | ACD company | #805641 |
| ADAM12 probe | ACD company | #432561 |
| **Biological samples** | | |
| Human wound samples | Karolinska Institutet Biobank | |
| Human plastic surgery samples | Karolinska Institutet Biobank | |
| Human venous ulcer samples | Dermatology Hospital of Chinese Academy of Medical Sciences | |
| Mouse wound samples (WT and DB) | Cyagen Biosciences | |
| **Chemicals, peptides, and recombinant proteins** | | |
| Dynabeads™ MyOne™ Streptavidin C1 | Thermo Fisher | 65001 |
| Pierce™ Protein A/G Magnetic Beads | Thermo Fisher | 88802 |
| Pierce™ 16% Formaldehyde (w/v), Methanol-free | Thermo Fisher | 28906 |
| Glycine | Sigma-Aldrich | 50046 |
| ProLong™ Diamond Antifade Mountant with DAPI | Thermo Fisher | P36966 |
| TRIzol™ Reagent | Thermo Fisher | 15596018 |
| Red Blood Cell Lysis Solution | Miltenyi Biotec | 130-094-183 |
| EpiLife™ Medium, with 60 µM calcium | Thermo Fisher | MEPI500CA |

| | | |
|---|---|---|
| Human Keratinocyte Growth Supplement (HKGS) | Thermo Fisher | S0015 |
| DMEM, high glucose | Thermo Fisher | 11965092 |
| TritonX100 | Merck | X100-100ML |
| Dispase II, powder | Thermo Fisher | 17105041 |
| Recombinant Human TNF-alpha Protein | R&D Systems | 210-TA-020 |
| BSA | Merck | 10711454001 |
| Pierce™ IP Lysis Buffer | Thermo Fisher | 87787 |
| Recombinant Proteinase K Solution (20 mg/mL) | Thermo Fisher | AM2546 |
| 2x Laemmli Sample Buffer | Bio-Rad Laboratories | 1610737EDU |
| AREG (20ng/mL) | Novoprotein | CG04 |
| TGFalpha (20ng/mL) | SinoBiological | 11252-HNAE |
| HB-EGF (20ng/mL) | SinoBiological | 10325-HNAB |
| HGF (20ng/mL) | Novoprotein | CJ72 |
| ANGPTL4 (500ng/mL) | Novoprotein | CW53 |
| EREG (20ng/mL) | GenScript | Z02865 |
| APRIL (TNFSF13) (500ng/mL) | Novoprotein | Cat. No.:CU89 |
| CXCL6 (20ng/mL) | Novoprotein | C598 |
| ADAM17 (50ng/mL) | Proteintech | Ag32418 |
| LTB (20ng/mL) | Abmart | RKRB5609S |
| CXCL1 (20ng/mL) | Novoprotein | C597 |
| CXCL5 (20ng/mL) | Novoprotein | CF14 |
| IL1B (20ng/mL) | Novoprotein | CG93 |
| **Critical commercial assays** | | |
| Whole Skin Dissociation Kit, human | Miltenyi Biotec | 130-101-540 |
| Dead Cell Removal Kit | Miltenyi Biotec | 130-090-101 |
| Pierce™ RNA 3' End Desthiobiotinylation Kit | Thermo Fisher | 20163 |
| Chromium Next GEM Single Cell 3′ Reagent Kits v3.1, 16 rxns | 10X Genomics | PN-1000268 |
| Chromium Next GEM Chip G Single Cell Kit, 48 rxns | 10X Genomics | PN-1000120 |
| Dual Index Kit TT Set A | 10X Genomics | PN-1000215 |
| RNAscope® Multiplex Fluorescent Reagent Kit v2 | Advanced Cell Diagnostics, Inc | 323100 |
| Visium Spatial Gene Expression Slide & Reagents Kit | 10X Genomics | 1000187 |
| Visium Spatial Gene Expression Starter Kit | 10X Genomics | 1000200 |
| Visium Spatial Tissue Optimization Slide & Reagents Kit | 10X Genomics | 1000193 |

| | | |
|---|---|---|
| RevertAid First Strand cDNA Synthesis Kit | Thermo Fisher | K1622 |
| Power SYBR™ Green PCR Master Mix | Thermo Fisher | 4368708 |
| TaqMan™ Universal PCR Master Mix | Thermo Fisher | 4304437 |
| **Deposited data** | | |
| Human skin acute wound single-cell RNA-seq data | This paper | GEO: GSE241132 |
| Human venous ulcer single-cell RNA-seq data | This paper | GEO: GSE265972 |
| Human skin acute wound spatial transcriptomics data | This paper | GEO: GSE241124 |
| Mouse wound data (UW + DPW3) | This paper | GEO: GSE218430 |
| Single-cell RNA-seq datasets from human healthy adult skin and inflamed skin diseases | Reynolds et al., 2021[1] | E-MATB-8142 |
| Diabetic foot ulcers single-cell RNA-seq data | Theocharidis et al., 2022[2] | GEO: GSE165816 |
| Human acute wound skin bulk RNA-seq data | Liu et al., 2022[3] | GEO: GSE174661 |
| Psoriatic skin single-cell RNA-seq data | Ma et al., 2023[4] | GEO: GSE173706 |
| **Experimental models: Cell lines** | | |
| Human Epidermal Keratinocytes, adult (HEKa) | Thermo Fisher | C0055C |
| **Oligonucleotides** | | |
| **qPCR primers** | | |
| FOSL1 | Sangon | Forward: TGACCACACCCTCCCT AACTC Reverse: CTGCTGCTACTCTTGCG ATGA |
| CXCL1 | IDT | Hs.PT.58.39039397 |
| GAPDH | IDT | Hs.PT.39a.22214836 |
| **siRNAs** | | |

| | | |
|---|---|---|
| siFOSL1 | GenePharma | Sense: GAGGGCAGCUGCUAUUUAUTT Antisense: AUAAAUAGCAGCUGCCCUCTT |
| **Software and algorithms** | | |
| CellRanger v5.0.1 | 10X genomics | https://www.10xgenomics.com/support/single-cell-gene-expression |
| Spaceranger v1.2.0 | 10X genomics | https://www.10xgenomics.com/support/software/space-ranger |
| R v4.1.1 and v4.2.3 | R Core Team | https://www.r-project.org/ |
| RStudio | Posit | https://posit.co/download/rstudio-desktop/ |
| Seurat v4 | Hao et al., 2021[5] | https://github.com/satijalab/seurat/ |
| Scrublet v0.2.3 | Wolock et al., 2019[6] | https://github.com/swolock/scrublet |
| DoubletFinder v2.0.3 | McGinnis et al., 2019[7] | https://github.com/chris-mcginnis-ucsf/DoubletFinder |
| Harmony v0.1 | Korsunsky et al., 2019[8] | https://github.com/immunogenomics/harmony |
| Canonical correlation analysis (CCA) | Butler et al., 2018[9] | https://github.com/satijalab/seurat/ |
| MiloR v1.6.0 | Dann et al., 2022[10] | https://github.com/MarioniLab/miloR |
| pySCENIC v0.11.2 | Aibar et al., 2017[11] | https://scenic.aertslab.org/ |
| CellOracle v0.10.14 | Kamimoto et al., 2023[12] | https://github.com/morris-lab/CellOracle |
| scikit-learn python package (Gaussian Mixture Model) | https://scikit-learn.org/stable/ | https://scikit-learn.org/stable/ |
| Monocle3 | Cao et al., 2019[13] | https://cole-trapnell-lab.github.io/monocle3 |
| CellRank v1 | Lange et al., 2022[14] | https://cellrank.org |
| clusterProfiler v4 | Wu et al., 2021[15] | https://github.com/YuLab-SMU/clusterProfiler |

| | | |
|---|---|---|
| MultiNicheNet | Browaeys et al., 2023[16] | https://github.com/saeyslab/multinichenetr |
| CellChat (v1.4.0) | Jin et al., 2021[17] | https://github.com/sqjin/CellChat |
| Cell2location | Kleshchevnikovet al., 2022[18] | https://github.com/BayraktarLab/cell2location/ |
| AutoGeneS | Aliee et al., 2021[19] | https://github.com/theislab/AutoGeneS |
| Cytoscape v3.8.2 | Shannon et al., 2003[20] | https://cytoscape.org/ |
| Image J | NIH | https://imagej.nih.gov/ij |
| BioRender | https://www.biorender.com | https://www.biorender.com |

### RESOURCE AVAILABILITY

#### Lead contact

#### Materials availability

All unique materials and reagents generated in this study are available from the Lead contact with a completed material transfer agreement.

#### Data and code availability

Original data sequenced in this study have been deposited in gene expression omnibus (GEO) under assession numbers: GSE241132, GSE265972, GSE241124, and GSE218430. All original code has been deposited at GitHub https://github.com/Zhuang-Bio/scRNA_STseq_human_wounds_paper_scripts. The single-cell and spatial gene expression data can be explored at the web porter https://www.xulandenlab.com/tools. Any additional information required to reanalyze the data reported in this paper is available from the lead contact upon request.

### EXPERIMENTAL MODEL AND STUDY PARTICIPANT DETAILS

#### Human wound sample collection

Five healthy volunteers were enrolled at the Karolinska University Hospital, Stockholm, Sweden. Three full-thickness skin wounds were created on the upper buttock area of each donor using a 4-mm biopsy punch. Wound edge tissues were collected on days 1, 7, and 30 using a 6-mm biopsy punch. The samples were transferred to the laboratory for single-cell RNA sequencing (10X Genomics) or snap-frozen for spatial transcriptomics (10X Genomics). Written informed consent was obtained from all donors for the collection and use of tissues for research. This study was approved by the Stockholm Regional Ethics Committee and conducted according to the Declaration of Helsinki's principles.

Venous ulcer samples and age- and body-location-matched healthy control samples from nine donors were collected at the Dermatology Hospital of Chinese Academy of Medical Sciences, Nanjing, China. Patients with apparent soft tissue infections or requiring systemic antibiotic treatment were excluded. After obtaining written informed consent from the patients, chronic wound edge tissue samples were collected using a 4-mm biopsy punch following a local lidocaine injection. This study was approved by the Ethics Committee of Institute of Dermatology, Chinese Academy of Medical Sciences (Ethic permission number: 2021-KY-059).

### METHOD DETAILS

#### Single-cell RNA sequencing

Wound skin tissues were incubated with 5U/ml Dispase II at 4°C overnight. The epidermis was gently separated from the dermis using tweezers and incubated in 0.025% Trypsin/ EDTA (Thermo Fisher) for 10-15 minutes at 37 °C. The dermal cell suspension was prepared using the whole skin dissociation kit (Miltenyi Biotec) following the manufacturer's instructions. To capture as many cell types as possible, equal amounts of epidermal and dermal cells were mixed. The red blood cells and dead cells were removed using the Red Blood Cell Removal Solution (Miltenyi Biotec) and Dead Cell Removal Kit (Miltenyi Biotec), respectively. After cleaning, viable cells were loaded onto the Chromium Controller (10X Genomics). Samples were processed for single-cell encapsulation and cDNA library generation using the Chromium Next GEM Single Cell 3′ Reagent Kits v3.1 (10× Genomics). The libraries were sequenced on an Illumina Novaseq6000 platform to achieve an average of approximately 50K read-pairs per cell.

#### Spatial transcriptomic sequencing

Fresh frozen skin and acute wound tissues were embedded in the Optimal Cutting Temperature compound (OCT, Sakura Tissue-TEK) on dry ice. Sections were fixed in methanol and imaged after hematoxylin and eosin (H&E) staining to assess the morphology and quality of the tissues. The optimal permeabilization time for wound sections was determined to be 15 minutes following the manufacturer's instructions (10x Genomics, Visium Spatial Tissue Optimization). Spatial gene expression libraries from 16 wound skin sections were then generated according to the instructions of the Visium Spatial Gene Expression Kit from 10X Genomics. The libraries were sequenced using the Illumina NovaSeq6000 platform to generate approximately 150 M read-pairs per section.

**scRNA-seq data processing**

The single-cell sequence data were mapped with CellRanger (version 5.0.1) to a manually built human reference genome GRCh38 with the annotation file GENCODE version 38. The raw gene expression matrix contained 65,462 cells and 27,973 genes. The low-quality cells expressing <500 genes, >20% mitochondrial genes, and <1000 gene counts were filtered out. Mitochondrial and hemoglobin genes, as well as genes expressed in fewer than ten cells, were excluded. Potential doublets were detected using Scrublet[6] (v0.2.3) and DoubletFinder[7] (v2.0.3). Cells identified as doublets by both tools or those in clusters with multiple distinct cell type markers were excluded. In total, 58,823 cells and 25,778 genes from 12 samples of acute wounds were retained for downstream analysis.

Data normalization and scaling were performed using the SCTransform[21] package, regressing out mitochondrial percentage and cell cycle effects. Cell cycle analysis was conducted using the CellCycleScoring function based on the normalized gene expression. The top 4000 variable genes were used for principal component analysis (PCA). During data integration, sample-to-sample batch effects were corrected using top 40 PCs as input for the RunHarmony[8] function. Uniform Manifold Approximation and Projection (UMAP) and k-nearest neighbors graph were generated using the RunUMAP and FindNeighbors functions in Seurat[5] (v4), respectively. Major cell clusters were identified using the Louvain graph-based algorithm with a resolution of 0.8, resulting in 27 clusters. Differentially expressed genes among clusters were calculated using the FindAllMarkers functions with the MAST method. Genes with adjusted *P*value<0.05, log fold change>0.25, and detected in at least 25 percent of cells were considered significantly high in the cluster. Clusters were annotated based on each cluster's top marker genes ranked by fold changes and well-documented signature genes of distinct cell types.

Before sub-clustering analysis, cells of low-quality keratinocyte cluster Bas-II from major clusters were filtered out. Subpopulation analyses of keratinocytes, fibroblasts, angiogenic cells, myeloid cells, and lymphoid cells were performed individually using the same pipeline as for major cluster identification, including normalization, variable feature selection, batch correction, dimensionality reduction, and unsupervised clustering, but with a resolution of 0.5.

Neutrophil analysis was performed using unfiltered count matrices of samples after running CellRanger. Cells with fewer than 100 expressed genes were filtered out, retaining only those not included in the above analysis. Initial neutrophils were selected based on the expression profiles of well-known markers (FCGR3B, CMTM2, CXCR2, PROK2, LINC01506)[22]. Cells were further filtered after clustering analysis based on the neutrophil scores of each cluster. The refined neutrophils were extracted and re-ran the normalization, scaling, and clustering steps with a resolution of 0.3.

**Differential abundance testing with Milo**

We tested for differential cell-state abundances of subpopulations across wound healing using the MiloR package (v1.6.0)[10]. Specifically, a K-nearest neighbors (KNN) graph was built using the graph 'HARMONY' slot from the adjacency matrix of the processed Seurat object with the parameters: k = 40 and d=30. Cells were assigned to the neighborhoods based on the KNN graph using the 'makeNhoods' function (prop=0.1). To explore variations in cell counts between neighboring wound healing points (pairwise comparisons), cells from each sample in each neighborhood were counted. Differential neighborhood abundance testing was performed using a generalized linear model (GLM). Differentially abundant cell neighborhoods with SpatialFDR ≤ 0.1 were plotted using the 'plotNhoodGraphDA' function.

**SCENIC and CellOracle analyses**

The pySCENIC (v0.11.2)[11] was utilized to investigate the role of transcriptional regulators in human skin wound healing, following the package's tutorial. Raw count expression matrices of cell types were first prepared to construct co-expression modules between transcription factors (TFs hg38) and potential target genes ranked by importance. Modules showing significant motif enrichment remained, and TFs with directed targets in these modules were defined as regulons. Each regulon was then assigned an activity score using the AUcell function. The top 5 regulons for each cell type were highlighted based on the scaled activity Z-scores across other cell types. The regulon network was visualized using Cytoscape[20] (v3.8.2) software.

*In silico* TF perturbation of gene regulatory networks (GRNs) was performed using CellOracle (v0.10.14)[12] package. Based on a pre-built GRN of human (hg38) from curated transposase-accessible chromatin with sequencing (ATAC-seq) data, we simulated cell identity shifts in response to TF FOSL1 knockout and overexpression, setting the expression value to 0 and 2, respectively. The simulated overexpression value exceeded the detected gene expression. Subsequently, we compared the simulated TF perturbation vector field with the natural development vector field by calculating the perturbation score (PS). Positive and negative PSs denoted the promotion and inhibition of cell differentiation, respectively.

**Trajectory analysis of keratinocytes and fibroblasts**

To infer the differences in epidermal cell trajectories between intact skin and wound conditions, we separated all keratinocytes into wound and non-wound cells using a Gaussian mixture model (GMM) in the 'scikit-learn' Python package[23]. We first filtered out cells that did not express KRT6A, a strong marker of wound-induced cells[24,25]. The positive cells were categorized into '0' and '1' classes, representing low- ($KRT6A^{dim}$) and high-expressing ($KRT6A^{+}$) cells. The wound cells comprised $KRT6A^{+}$ cells, while the rest of the keratinocytes ($KRT6A^{dim}/KRT6A^{-}$) were defined as non-wound cells. Pseudotime trajectory analysis of wound and non-wound keratinocytes was performed using Monocle3[13] package. The basal cell cluster (Bas-I) was selected as a starting point of the pseudotime trajectory. The differentially expressed driving genes along the trajectory were determined using Moran's I test in the 'graph_test' function, with the filtering criteria: q_value<0.00001 and morans_I>0.25.

RNA velocity analysis of fibroblasts was carried out using the CellRank (version 1)[14] package, which predicted the cell differentiation trajectory and its directionality based on the spliced and unspliced mRNA content. The initial and terminal states were identified using a deterministic mode in 'cr.tl.initial_states' and 'cr.tl.terminal_states' functions.

**Gene ontology analysis**

Gene ontology (GO) analysis of all gene clusters was computed using the Fisher exact test in the clusterProfiler[15] package. GO biological process (BP) terms were filtered by the adjusted *P*value<0.05 and enriched gene count>5.

**M1 and M2 signature**

Pro-inflammatory macrophage (M1) and anti-inflammatory macrophage (M2) signatures were derived from Table 1 published by Martinez et al. (2006)[26]. Those in common with top marker genes of macrophage clusters were used to calculate the M1- and M2-like scores using the 'AddModuleScore' function in the Seurat package[5].

**Cell-to-cell communication analysis**

To study putative cell-cell interactions across the wound healing process, we used the MultiNicheNet[16] R package to infer the ligand-receptor pairs in the acute wound scRNA-seq dataset. The MutiNicheNet is a novel framework based on prior knowledge of ligand-receptor and ligand-target networks (version 2) that better explores cell-cell communications from multi-sample, multi-condition scRNA-seq data. Significant ligand-receptor pairs between cell types were determined by the high expression of each pair, as well as differentially expressed target genes of ligands in receiving cell types of different conditions, using the 'multi_nichenet_analysis' function with criteria: min_cells=10, logFC_threshold=0.50, p_val_threshold=0.05, fraction_cutoff=0.05, top_n_tatget=250.

In addition, we applied the CellChat[17] package to perform differential signaling changes within basal and spinous migrating keratinocytes between intact skin and wound conditions. Cells from each condition were imported into the CellChat analysis individually. The major signaling contributors of each cell type were calculated based on signaling network likelihoods using the 'netAnalysis_computeCentrality' function with a threshold of p-value<0.05. The differential outgoing and incoming interaction strengths of each cell population in the cell-cell communication network between the two conditions were computed using the 'netAnalysis_signalingRole_scatter' function.

**ST-seq data processing**

Spatial sequencing data of human acute wounds was mapped to the GRCh38 human genome using Space Ranger (v1.2). Spots with less than 100 genes expressed and in low-quality clusters were excluded. We also filtered out *MALAT1*, mitochondrial, and hemoglobin-related genes. In total, 22,915 spots and 36,578 genes from 16 sections remained. The data normalization and batch correction were performed using SCTransform[21] and Harmony[8], respectively. Dimensionality reduction, clustering (res=0.5), UMAP, and differentially expressed genes (DEGs) were carried out using the Seurat (v4)[5] package. Spot clusters were annotated based on DEGs and markers of distinct cell types from scRNA-seq. Spatial cell type distribution and gene expression were visualized using the 'SpatialDimPlot' and 'SpatialFeaturePlot' functions, respectively.

**Deconvolution of ST-seq data and wound bulk RNA-seq data**

To spatially map wound cell states defined by scRNA-seq data profiles in the Visum data, we used the Cell2location[18] package. In brief, we trained a negative binomial regression model to estimate reference transcriptomic signatures based on each cell type's top 100 marker genes profiled by scRNA-seq. We estimated the abundance of every cell type in each Visum spot using the inferred reference cell type signatures by decomposing spot mRNA counts. All parameters were set to default except for two: 1) the expected cell abundance (N_cells_per_location=20) determined by approximately counting the average numbers of nuclei of each spot in H&E images, and 2) regularisation of per-location normalization (detection_alpha = 20) to account for large variations in RNA detection sensitivity across different spots on Visium slides. The posterior distribution of cell abundance for each cell type in each spot was summarized as 5% quantile, representing high confidence, which was used for visualization and colocalization analysis. To identify microenvironments of spatial co-occurrence of cell types, we performed a non-negative matrix factorization (NMF) analysis of the high-confidence cell type abundances, setting the number of factors to R=15. A cell type was considered localized in a microenvironment if its fraction was over 0.1.

Bulk RNA sequencing data of intact skin and wounds (Day 1 and Day 7 post-wounding) from our previous study[3] (GSE174661) was deconvoluted using the AutoGeneS[19] package. Centroids of cell types were first constructed from our scRNA-seq data of acute wounds using the top 4000 highly variable genes and distinct makers of each cell type. The bulk data was then deconvoluted based on these centroids using a regression method of Nu-support vector machine (Nu-SVR).

**Venous ulcer scRNA-seq analysis**

scRNA-seq data containing 48,346 high-quality cells from venous ulcer (VU) patients (n=4) and matched healthy controls (n=5) was analyzed using the same workflow and criteria as for acute wound scRNA-seq.

**Integration of wounded skin scRNA-seq datasets**

We retrieved public scRNA-seq datasets of healthy adult and inflamed disease skins from *Reynolds et al.* paper[1] (2021, E-MTAB-8142 psoriatic and eczematous skin) and Ma et al. paper[4] (2023 GSE173706 psoriatic skin), as well as diabetic foot ulcer (DFU) skins from *Theocharidis et al.*[2] (2022, GSE165816). For the DFU scRNA-seq, we included 9 healthy controls and two subgroups of DFU patients: those who healed the ulcers (Healer, DFU_H n=7) and those who failed to heal within 12 weeks post-surgery (Non-healer, DFU_NH n=4). Data integration across different scRNA-seq datasets, including acute wounds, VU, DFU, human adult skin, psoriasis, and eczema skin, was carried out using the Harmony[8] algorithm, setting each sample as a group variable.

Cell label transfer was performed using the 'FindTransferAnchors' and 'MapQuery' functions in Seurat[5], setting the cell types generated from our acute wound dataset as a reference. The average predicted scores of each cell type were used to assess the correlation of cell types across different scRNA-seq datasets. Furthermore, we refined cell labels using integrated unsupervised clustering results, resolving ambiguous assignments by aligning them with the majority cell type in each population to enable a precise comparison of cellular heterogeneity between acute and chronic wounds.

**Cross-species comparison of human and mouse wound healing**

To compare human and mouse wound healing, we integrated scRNA-seq data of human and mouse acute wounds at the inflammatory phase (GSE218430) using canonical correlation analysis (CCA). Before integration, we sampled the same number of cells from each sample in both datasets. The overlapped homologous genes of humans and mice were kept for integration. CCA[9] in Seurat (v4)[5] was used to integrate the human and mouse scRNA-seq datasets using the top 3000 variable genes and each sample as an integration anchor. The dynamic changes of cellular proportions of human-mouse joint clusters were traced according to the original cell-type assignments using the Sankey diagram. Conserved markers of integrated clusters were identified using the Seurat 'FindConservedMarkers' function. Differential expression analysis between human and mouse migrating keratinocytes was performed using the MAST test in Seurat 'FindMarkers' function[5]. Functional enrichment of DEGs in humans and mice was carried out using clusterProfiler[15], and the top 6 significant BP terms (adjusted *P*value<0.05 and enriched gene count>5) were plotted. Cross-species comparison of migration and proliferation scores in integrated migrating and proliferating clusters was calculated based on the top 10 conserved marker genes across species using the Seurat 'AddModuleScore' function.

Human-specific coding and non-coding genes were retrieved from previous literature[27,28], and their expression was visualized using the Seurat 'DoHeatmap' function.

**Fluorescent in situ hybridization (FISH)**

Probes for MMP3, KRT6B, and ADAM12 (Hs-MMP3:#403421, Hs-KRT6B:#805641, Hs-ADAM12:#432561) were designed by Advanced Cell Diagnostics (ACD) in Silicon Valley, CA. Human skin and wound slides were prepared according to the manufacturer's instructions. After fixation and dehydration using 50%, 70%, and 100% ethanol, the slides were treated with Protease IV (ACD) and incubated at room temperature for 30 minutes. Subsequently, the slides were incubated with the probes for two hours at 40 °C, using the HybEZ™ II Hybridization System and the RNAscope® Multiplex Fluorescent Reagent Kit v2 (ACD). The hybridization signals were amplified per the manufacturer's instructions and captured using a Zeiss Axio Scan Z1 slide scanner.

**Immunofluorescence staining and microscopy**

Paraffin-embedded tissue sections were deparaffinized and rehydrated using xylene and a series of graded ethanol solutions. Antigen retrieval was performed in citric acid buffer (10 mM, pH 6.0). The sections were then blocked with 2.5% bovine serum albumin (BSA) in Tris-buffered saline with 0.1% Tween-20 (TBST). Next, the sections were incubated overnight at 4°C with primary antibodies specific to the anti-FOSL1 (1:100 dilution, Thermofisher, #PA5-40361), anti-IL1b (1:150 dilution, Abcam, ab156791), anti-CD68 (1:100 dilution, Conjugated with Alexa Fluor® 488, Abcam, ab222914). After primary antibody incubation, the sections were treated with Alexa Fluor 555 Donkey anti-Rabbit IgG (H+L) Highly Cross-Adsorbed secondary antibody (cat. A-31572, ThermoFisher Scientific) or Goat anti-Mouse IgG (H+L) Highly Cross-Adsorbed Secondary Antibody, Alexa Fluor Plus 647 (cat. A-32728, ThermoFisher Scientific) diluted at 1:1000 in TBST. To visualize cell nuclei, we counter-stained the sections with DAPI and mounted them using an anti-fading polyvinyl alcohol mounting medium (ThermoFisher Scientific). Immunofluorescence staining was observed using a Confocal fluorescence microscope.

**RNA extraction and qRT-PCR**

Total RNA was isolated from human *in vivo* wounds using the miRNeasy mini kit (Qiagen), followed by cDNA synthesis using the RevertAid First Strand cDNA Synthesis Kit (ThermoFisher

Scientific). Specific premixed primers and probes for CXCL1 and GAPDH were designed by Integrated DNA Technologies (IDT, Leuven, Belgium). Gene expression levels were quantified using TaqMan expression assays (ThermoFisher Scientific) and normalized to the housekeeping gene GAPDH. The comparative $2^{-\Delta\Delta CT}$ method was used for gene expression quantification, and all reactions were conducted on QuantStudio 6 or 7 platforms (Applied Biosystems, Waltham, MA).

**Keratinocyte culture, treatment, RNA extraction, and qRT-PCR**

Human adult primary keratinocytes (Lifeline® Cell Technology) were cultured in DermaLife Basal Medium supplement with DermaLife K LifeFactors® Kit and antibiotics [penicillin (100 U/ml), streptomycin (100 U/ml); Thermo Fisher Scientific] at 37°C, 5% CO2. For keratinocyte treatment, the cells were plated into 24-well plates and treated with growth factors or cytokines as described in the Key Resource Table when reaching about 70% confluency. After incubation with growth factors or cytokines for 24 hours, cells were lysed in RNAiso Plus (Takara,9109). RNA was isolated and reverse transcribed into complementary DNA using PrimeScript™ RT Master Mix (Takara, RR036A) according to the manufacturer's instructions. qRT–PCR was performed using TB Green® Premix Ex Taq™ II (Tli RNaseH Plus) (Takara, RR820A) on a Roche LightCycler®96 system. GAPDH was used as the internal control. Results were normalized to the internal control, and the comparative $2^{-\Delta\Delta CT}$ method was used to quantify gene expression.

**Cell migration assay**

Keratinocytes were plated into 6-well plates and treated with growth factors or cytokines as described in the Key Resouse Table when reaching about 70% confluency. After incubation with growth factors or cytokines for 24 hours, cells (95-100% confluency) were scratched using a 200ul pipette tip. The cells were cultured in a supplement-free medium and allowed to grow for 24 hours. Cell images at 0 h and 24 h were taken under a microscope at a magnification of × 10. The cell migration rate was analyzed by measuring the healed area of the scratch using the Image J software.

**Western blotting**

Primary keratinocytes were plated into 6-well plates and incubated for at least 24 hours. When they reached about 70% confluency, cells were treated with 20µM U0126 or DMSO for 30 minutes, and then 2 ng/mL CXCL1 was added and incubated for the indicated time. Cells were lysed by RIPA lysis buffer (P0013C, Beyotime) supplemented with protease and phosphatase inhibitors on ice for 10 minutes, and the cell debris was removed by centrifugation at 12,000 rpm at 4°C for 5 minutes. The lysate was boiled with SDS loading buffer, and equal amounts of protein were loaded onto 4-20% precast polyacrylamide gels (Tanon, 180-9110H) and then transferred onto nitrocellulose membranes (Pall Corporation, 66485). The membrane was blocked with 5% non-fat powdered milk in tris-buffered saline with tween-20 (TBST). After blocking, the membrane was incubated with primary antibodies (1:1000) at 4°C overnight, washed with TBST, and incubated with HRP-labeled goat anti-rabbit secondary antibody (1:2000) (Cell Signaling Technology, 7074). Protein bands were visualized using Clarity™ Western ECL Substrate (Bio-Rad Laboratories, 170-5061). The density of protein bands was quantified using ImageJ software. GAPDH served as the loading control.

**Softwares and statistics**

The tool used to visualize acute wound scRNA-seq was adapted from the R package ShinyCell[29]. The workflow and schematic summary of this study were created using BioRender. Statistical significances in migration assay and immunofluorescence staining quantification between groups were determined using either a two-tailed Student's t-test or ANOVA analysis facilitated by GraphPad Prism 8 (GraphPad Software Inc, California, USA). Cell proportion and gene expression were compared between groups using a quasi-binomial distribution model and Mann-Whitney U test in R, respectively. A significance threshold of $P<0.05$ was applied for all statistical tests. Data were presented as mean ± standard deviation (SD) or mean ± standard error of the mean (SEM).
