## Supplementary figures for "Spatiotemporal Single-Cell Roadmap of Human Skin Wound Healing"

**Figure S1 Analysis and integration of scRNA-seq datasets of human acute and chronic wounds related to Figure 1**

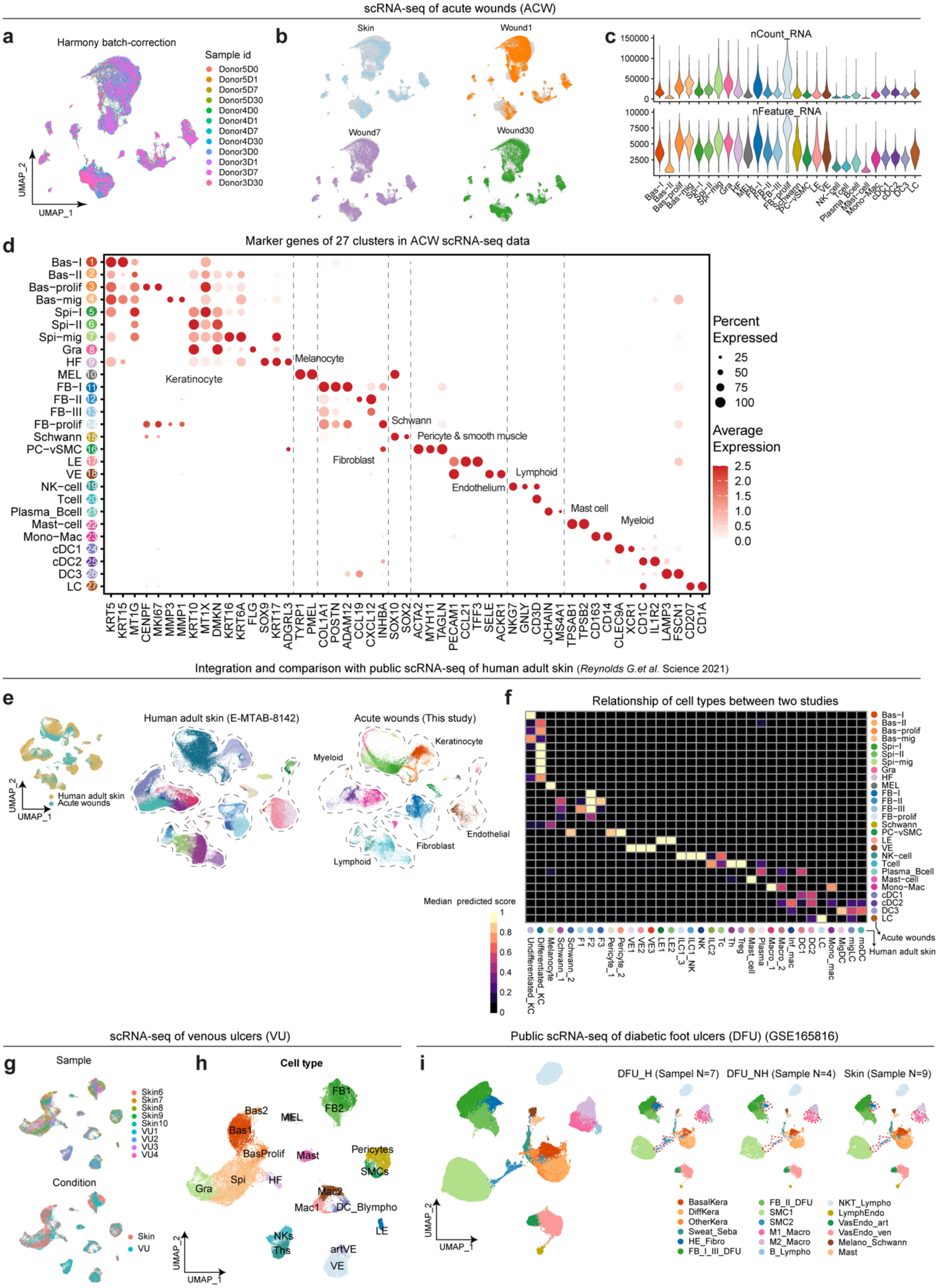

**Figure S1 Analysis and integration of scRNA-seq datasets of human acute and chronic wounds related to Figure 1.** UMAP showing the cell distribution of each sample (**a**) and condition (**b**) in scRNA-seq data of human skin and acute wounds after batch correction. (**c**) Violin plots showing each cluster's total read counts and feature gene counts in scRNA-seq data of human skin and acute wounds after quality control. (**d**) Dot plot showing the expression of distinctive markers for each cell population. Bas-/Spi-/Gra, basal/spinous/granular keratinocyte; -prolif/-mig, proliferating/migrating cells; HF, hair follicle; MEL, melanocyte; FB, fibroblast; PC-vSMC, pericyte, and vascular smooth muscle cell; LE/VE, lymphatic/vascular endothelial cell; NK-cell, natural killer cell; Th, T helper cell; Mono-mac, monocyte and macrophage; cDC, conventional dendritic cell; LC, Langerhans cell. (**e**) UMAP showing the integration of our scRNA-seq data of human acute wounds with a public scRNA-seq dataset of adult human skin (E-MTAB-8142). (**f**) Heatmap showing the median prediction scores of cell types between these two datasets using the TransferAnchors function in Seurat. F1/2/3, fibroblast 1/2/3; ILC, innate lymphoid cell; Tc, cytotoxic T cell; Treg, regulatory T cell; Inf., inflammatory; Mono mac, monocyte-derived macrophage; Mig., migratory; MoDC, monocyte-derived dendritic cell. Analysis and integration of scRNA-seq datasets of venous ulcer (VU) (**g**, **h**) and diabetic foot ulcer (DFU: GSE165816) (**i**). BasalKera/DiffKera: basal/differentiated keratinocytes; Sweat_Seba: sweat and sebaceous gland cells; HE-Fibro: healing enriched fibroblasts; _Macro: macrophages; B-Lympho: B-lymphocytes; NKT_Lympho: NK cells and T lymphocytes; LymphEndo: lymphatic endothelial cells; VasEndo_art/ven: arteriole/venule vascular endothelial cells; Melano_Schwann: melanocytes and Schwann cells.

**Figure S2 Analysis of acute wound spatial transcriptomic sequencing (ST-seq) data related to Figure 1d-f**

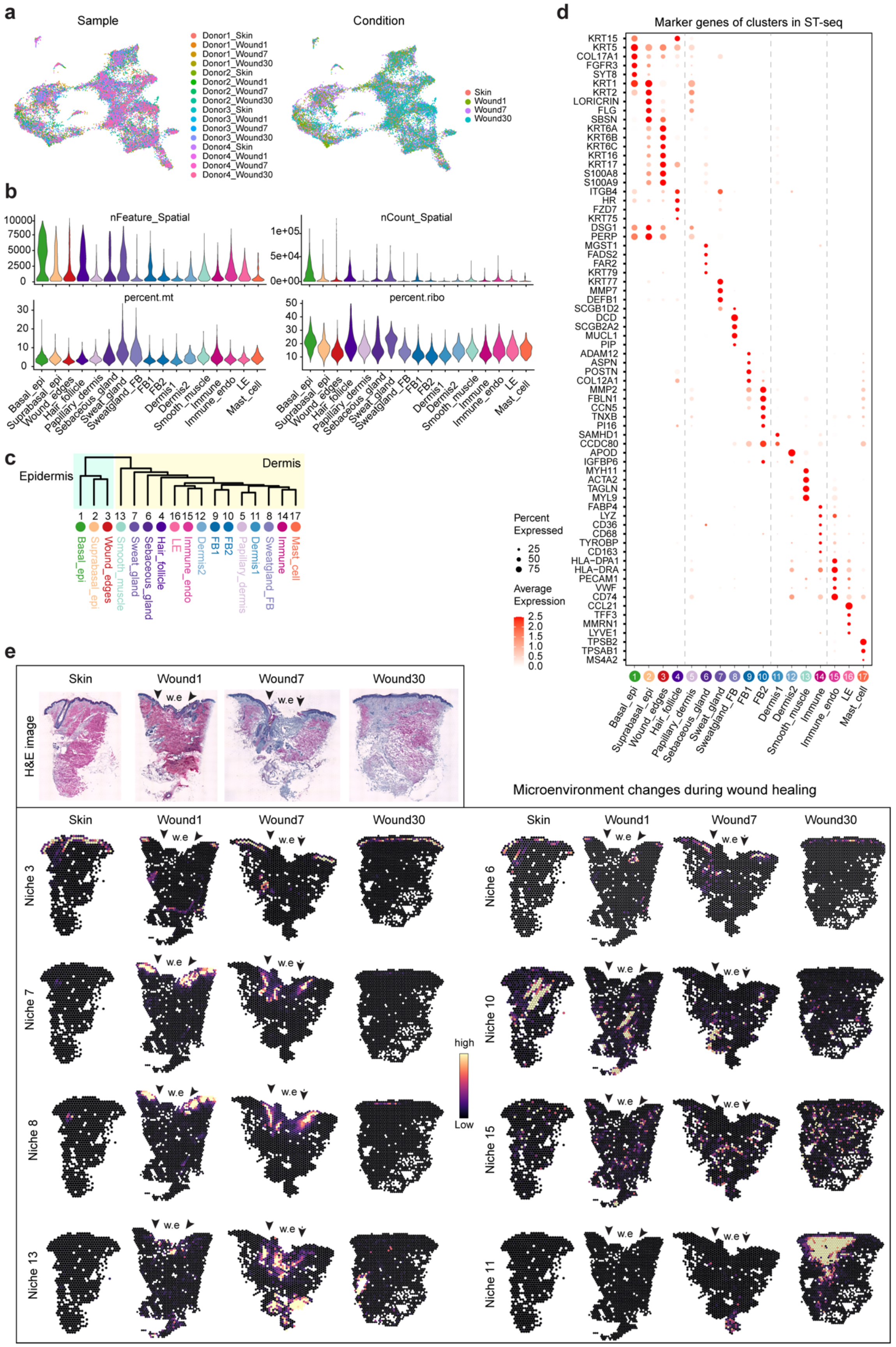

**Figure S2 Analysis of acute wound spatial transcriptomic sequencing (ST-seq) data related to Figure 1d-f** (**a**) UMAP showing the spot distribution of each sample and condition in ST-seq after batch correction. (**b**) Violin plots showing total feature counts, read counts, and the percentage of mitochondria and ribosome genes of each cluster in acute wound ST-seq data after quality control. (**c**) Unsupervised clustering dendrogram illustrating the relatedness of spot clusters based on a distance matrix constructed in gene expression space. (**d**) Dot plot showing the expression of distinctive markers for each population. epi, epidermis; FB, fibroblast; endo, endothelium; LE, lymphatic endothelium. (**e**) Spatial feature plots showing the changes of cell-type co-occurrence across wound healing. These niches were identified using non-negative matrix factorization (NMF), as shown in **Figure 1f**. w.e: wound edge. The hematoxylin and eosin (H&E) staining image of human skin and acute wounds from donor 4 is shown.

**Figure S3 Sub-clustering analysis of human wound keratinocytes related to Figures 2 and 3**

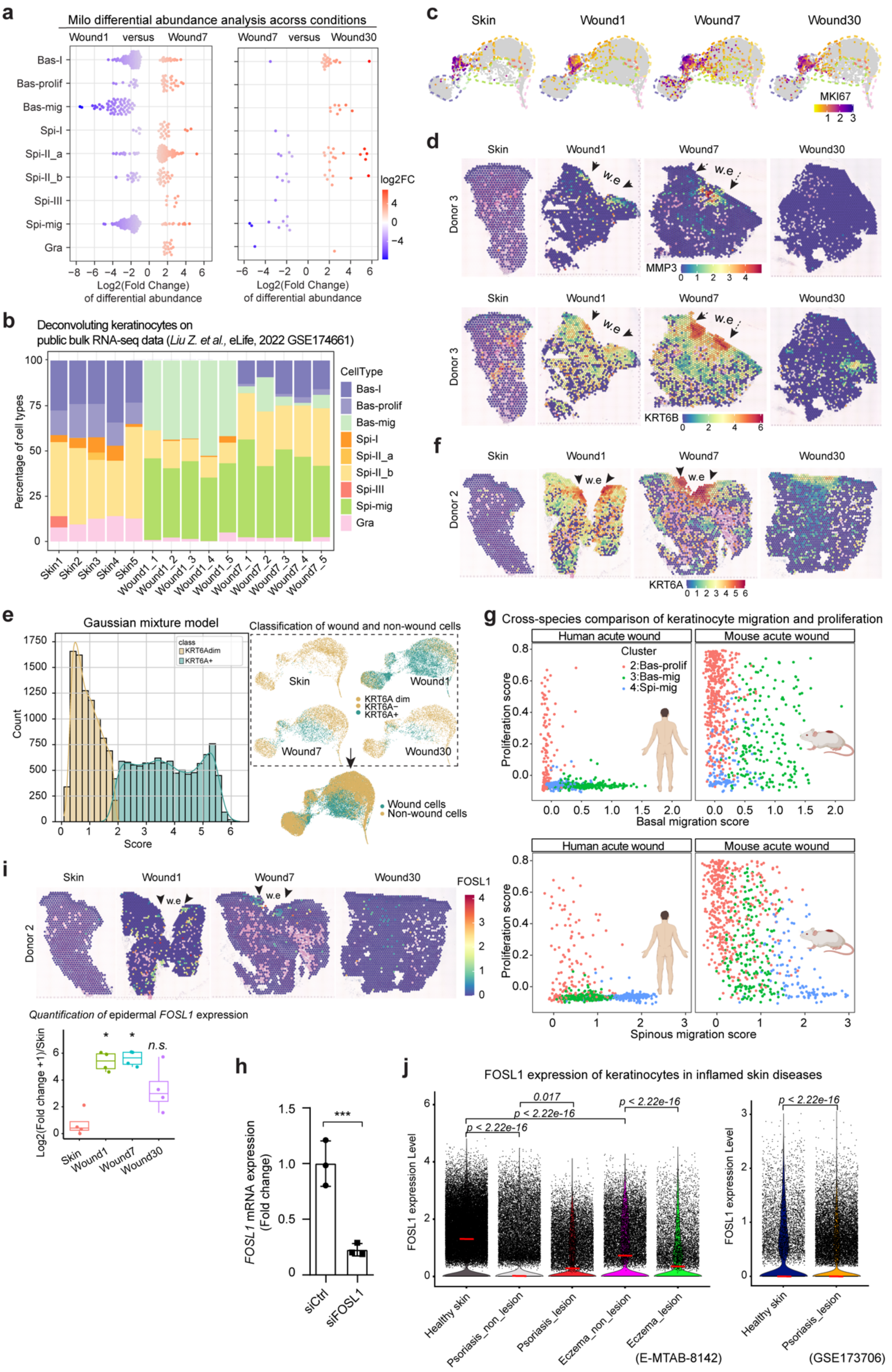

**Figure S3 Sub-clustering analysis of human wound keratinocytes related to Figures 2 and 3.** (**a**) Milo beeswarm plot showing the differential abundance of keratinocyte cell types between Wound7 and Wound1 (left panel), and Wound30 vs. Wound7 (right panel). Blue and red dots indicate significantly (SpatialFDR < 0.1) decreased (logFC < 0) and increased (logFC > 0) cell abundance, respectively. Color intensity indicates the degree of significance for each neighborhood. (**b**) Cell proportions of keratinocyte sub-clusters in the deconvolution of public bulk RNA-seq data of human skin and acute wounds from 5 donors (GSE174661). (**c**) Feature plots showing keratinocyte expression of proliferating cell marker MKI67 during wound healing. (**d**) Spatial feature plots showing MMP3 and KRT6B expression in ST-seq data of human skin and acute wounds. Arrows indicate the wound edges (w.e). (**e**) Classification of wound and non-wound cells using the Gaussian mixture model. $KRT6A^{-}$ and $KRT6A^{dim}$ cells were defined as non-wound cells, while the $KRT6A^{+}$ cells were wound cells. (**f**) Spatial feature plots showing KRT6A expression in ST-seq data of human skin and acute wounds. (**g**) Scatter plot showing the basal/spinous migration and proliferation scores of migrating (integrated clusters3&4) and proliferating (integrated cluster2) keratinocyte clusters in human and mouse wounds. The integrated clusters are shown in Figure 7a. (**h**) qRT-PCR analysis of *FOSL1* mRNA expression in human primary keratinocytes transfected with FOSL1 or control siRNAs (n=3). (**i**) Spatial feature plots showing FOSL1 expression in ST-seq data of human skin and acute wounds. Epidermal FOSL1 expression across wound healing was quantified. (**j**) Violin plots showing FOSL1 expression in keratinocytes of inflamed skin diseases. Data were extracted from public scRNA-seq datasets of psoriasis and eczema: E-MTAB-8142 and GSE173706. Significance was determined using the student t-test (**h**) and the Mann-Whitney U test (**i, j**), comparing other conditions to normal skin/control group, *: $p < 0.05$, n.s.: no significance.

**Figure S4 Sub-clustering analysis of immune cells during wound healing related to Figure 4**

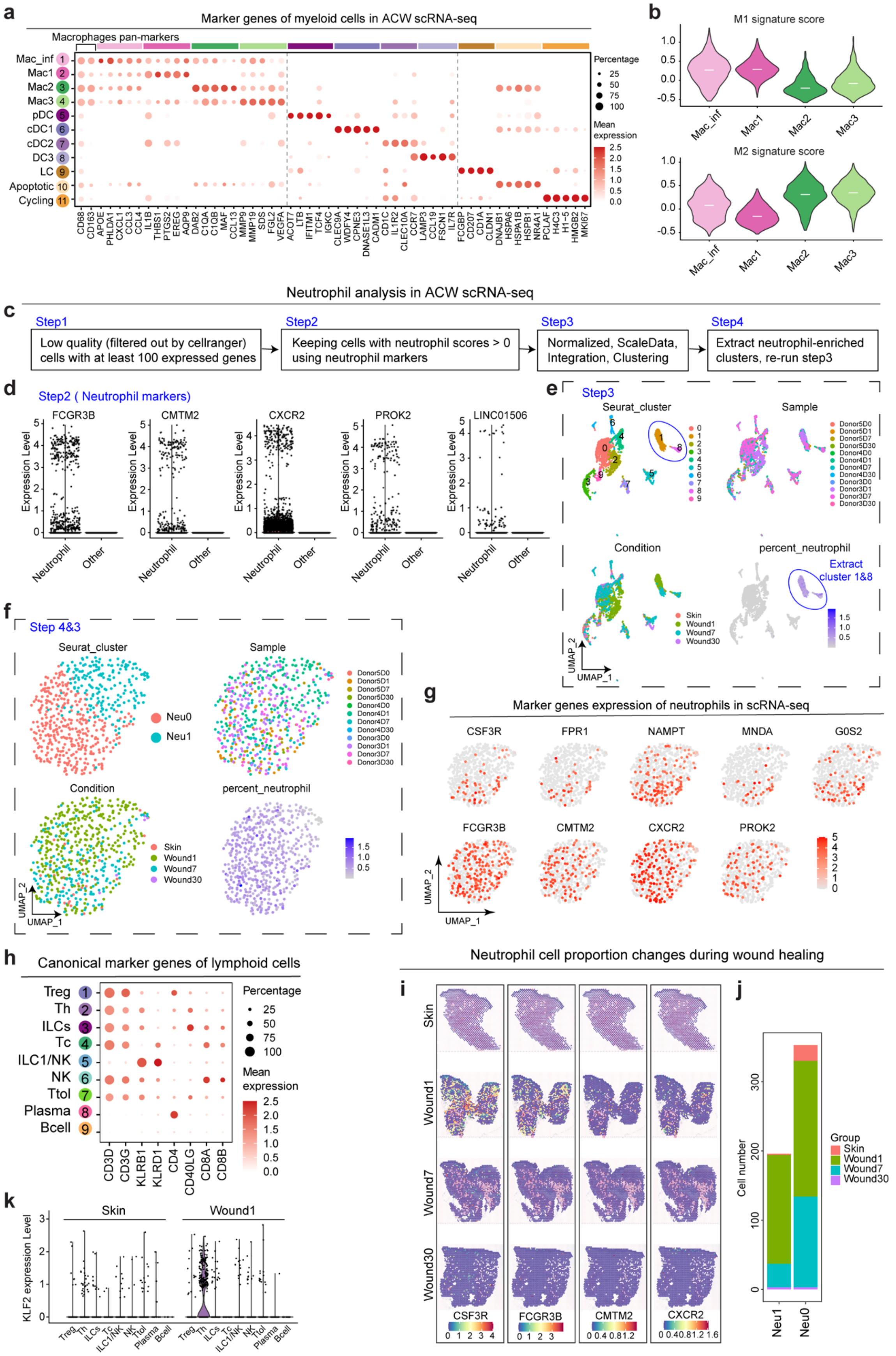

**Figure S4 Sub-clustering analysis of immune cells during wound healing related to Figure 4.** (**a**) Dot plot showing markers of myeloid cell clusters. Mac_inf, inflammatory macrophage; Mac1/2/3, macrophage 1/2/3; pDC, plasmacytoid dendritic cell; cDC1/2, conventional dendritic cell 1/2; LC, Langerhans cell. (**b**) Violin plots showing pro-inflammation (M1) and anti-inflammation (M2) gene signatures for the four macrophage clusters identified in human acute wounds. White lines indicate median scores. (**c-f**) Workflow of neutrophil clustering analysis. (**g**) Feature plots showing marker gene expression of neutrophils in scRNA-seq data of human skin and acute wounds. (**h**) Dot plot showing the expression of canonical markers of lymphoid cell clusters. Treg, regulatory T cell; Th, helper T cell; ILCs, innate lymphoid T cell; Tc, cytotoxic T cell; NK, natural killer T cell; Ttol, tolerant T cell. (**i**) Spatial feature plots showing neutrophil marker expression across wound healing. (**j**) Bar chart showing the proportional representation of neutrophils during wound healing. (**k**) Violin plot showing KLF2 expression in lymphoid cell types in Skin and Wound1 analyzed by scRNA-seq.

**Figure S5 Cell-to-cell crosstalk during human skin wound healing related to Figures 4 and 5**

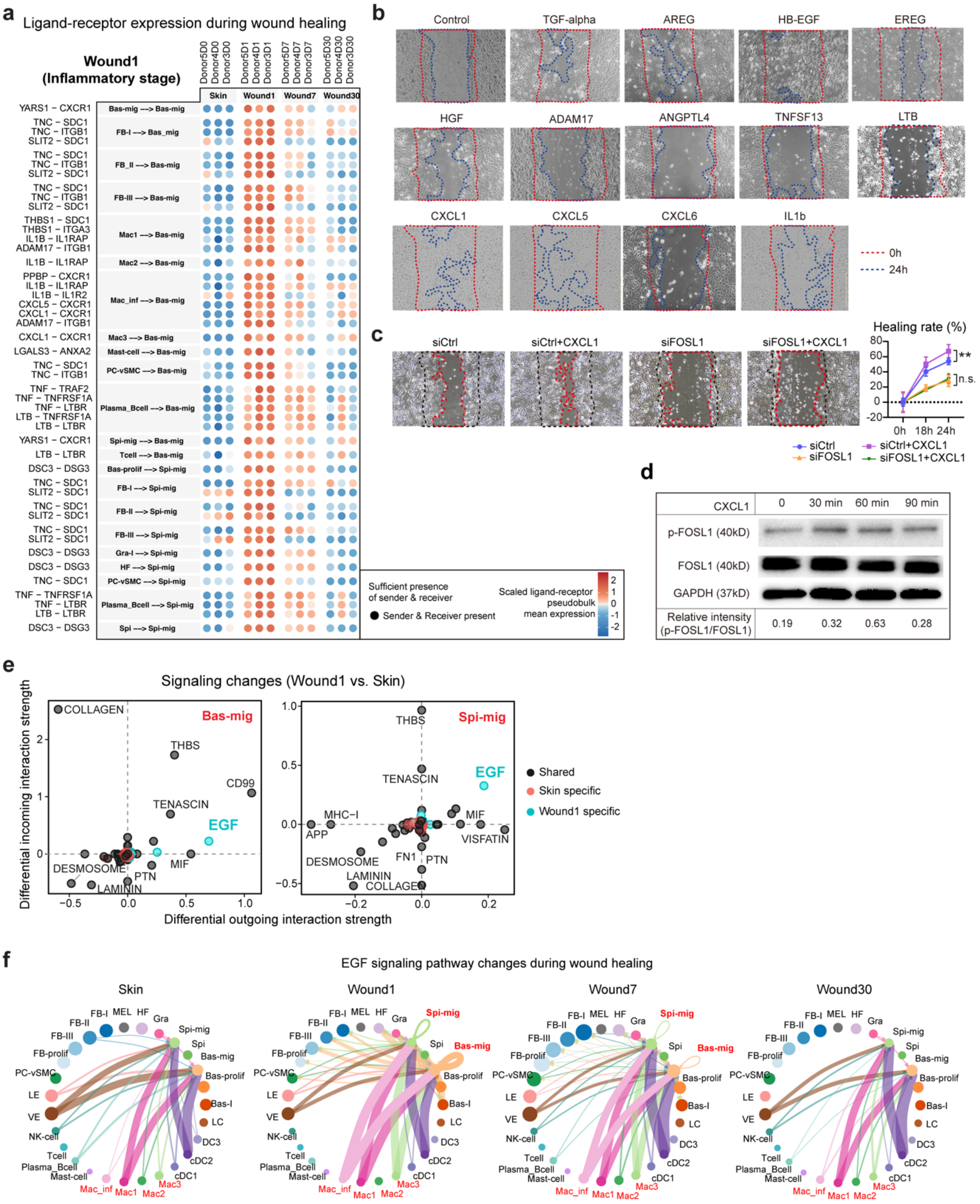


**Figure S5 Cell-to-cell crosstalk during human skin wound healing related to Figures 4 and 5.** (**a**) Heatmap showing the scaled average expression of the top 50 ligand-receptor pairs in Wound1. Each column represents a sample. (**b**) Scratch wound assay in primary human keratinocytes treated with growth factors or cytokines. (**c**) Scratch wound assay of primary human keratinocytes with or without

*FOSL1* expression silencing and treated with or without CXCL1. Significance was determined using a one-way ANOVA test, $^{**}P < 0.01$, n.s., no significance. (**d**) Western blot of phosphorylated and total FOSL1 proteins in keratinocytes treated with CXCL1 for 30-90 minutes. GAPDH was used as a loading control. (**e**) CellChat analysis of differential signals in basal and spinous migrating keratinocytes between Skin and Wound1. Blue and red dots represent Wound1- and skin-specific signals, respectively. (**f**) Circle plots showing dynamics of the EGF signaling pathway across skin wound healing. The color of the segments indicates the cell source of the signal. Segment size is proportional to the total interaction strength of each ligand-receptor pair in the corresponding cell types.

**Figure S6 Sub-clustering analysis of fibroblasts and angiogenic cells during wound healing related to Figure 5**

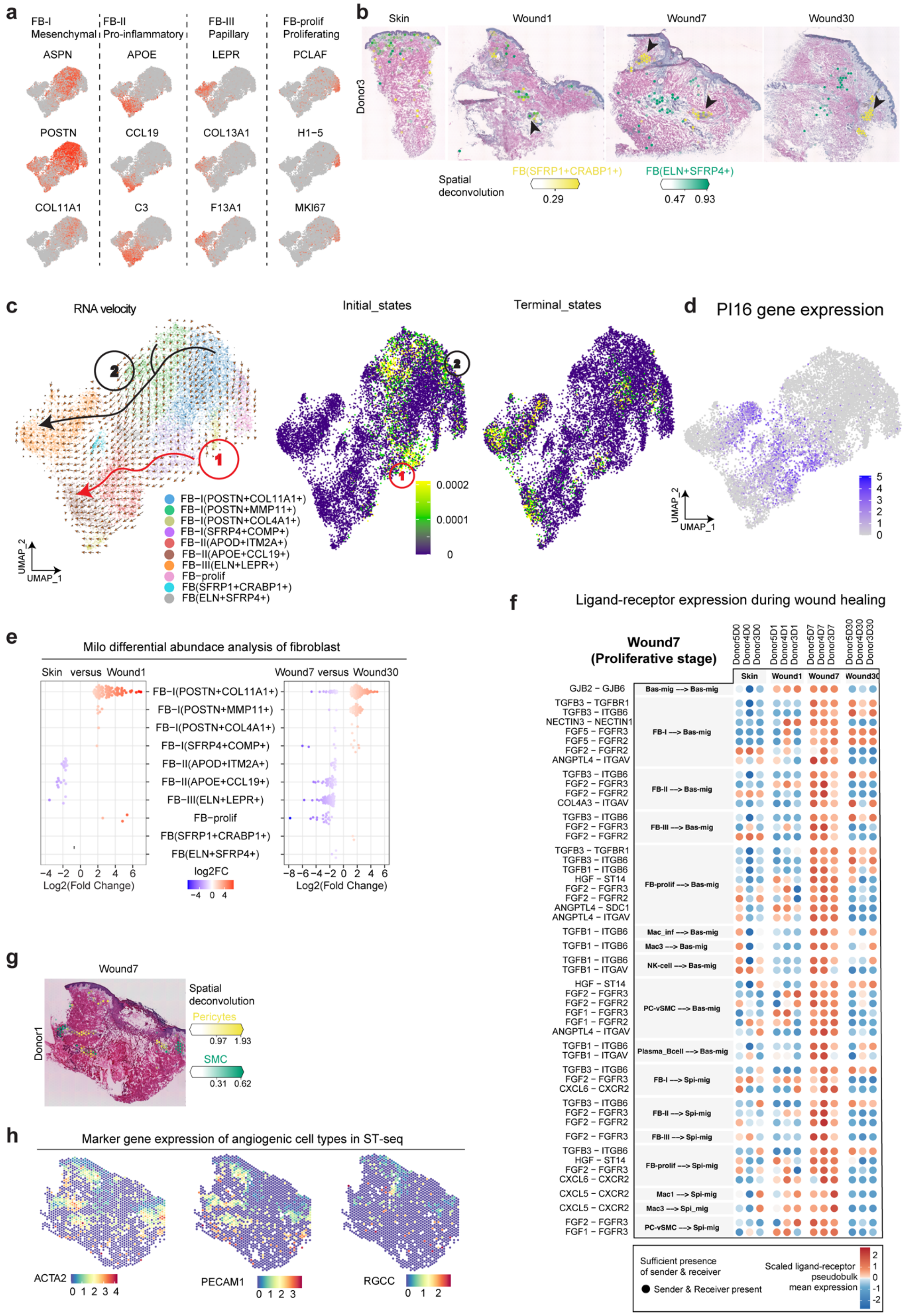

**Figure S6 Sub-clustering analysis of fibroblasts and angiogenic cells during wound healing related to Figure 5**. (**a**) Feature plots showing marker genes of reported fibroblast (FB) clusters. (**b**) Deconvolution of FB subclusters [FB($SFRP1^{+}CRABP1^{+}$), FB($ELN^{+}SFRP4^{+}$)] in ST-seq data of human skin and acute wounds. Arrows indicate hair follicles. (**c**) RNA velocity analysis showing two differentiation trajectories in FB clusters. (**d**) Feature plot showing progenitor FB marker PI16 expression in FB. (**e**) Milo beeswarm plot showing the differential abundance of FB cell types between Wound1 and Skin (left panel) and Wound30 vs. Wound7 (right panel). Blue and red dots indicate significantly (SpatialFDR < 0.1) decreased (logFC < 0) and increased (logFC > 0) cell abundance, respectively. Color intensity indicates the degree of significance for each neighborhood. (**f**) Heatmap showing the scaled average expression of the top 50 ligand-receptor pairs in Wound7. Each column represents a sample. (**g**) Deconvolution of pericytes and smooth muscle cells (SMC) in ST-seq data of Wound7. (**h**) Spatial feature plots showing angiogenic cell marker gene expression in Wound7. ACTA2 is a marker of pericytes and SMC. PECAM1 is a pan-marker of endothelium. RGCC is a marker of capillary vascular endothelium.

### Figure S7 Multi-facet pathological changes in chronic wounds related to Figure 6

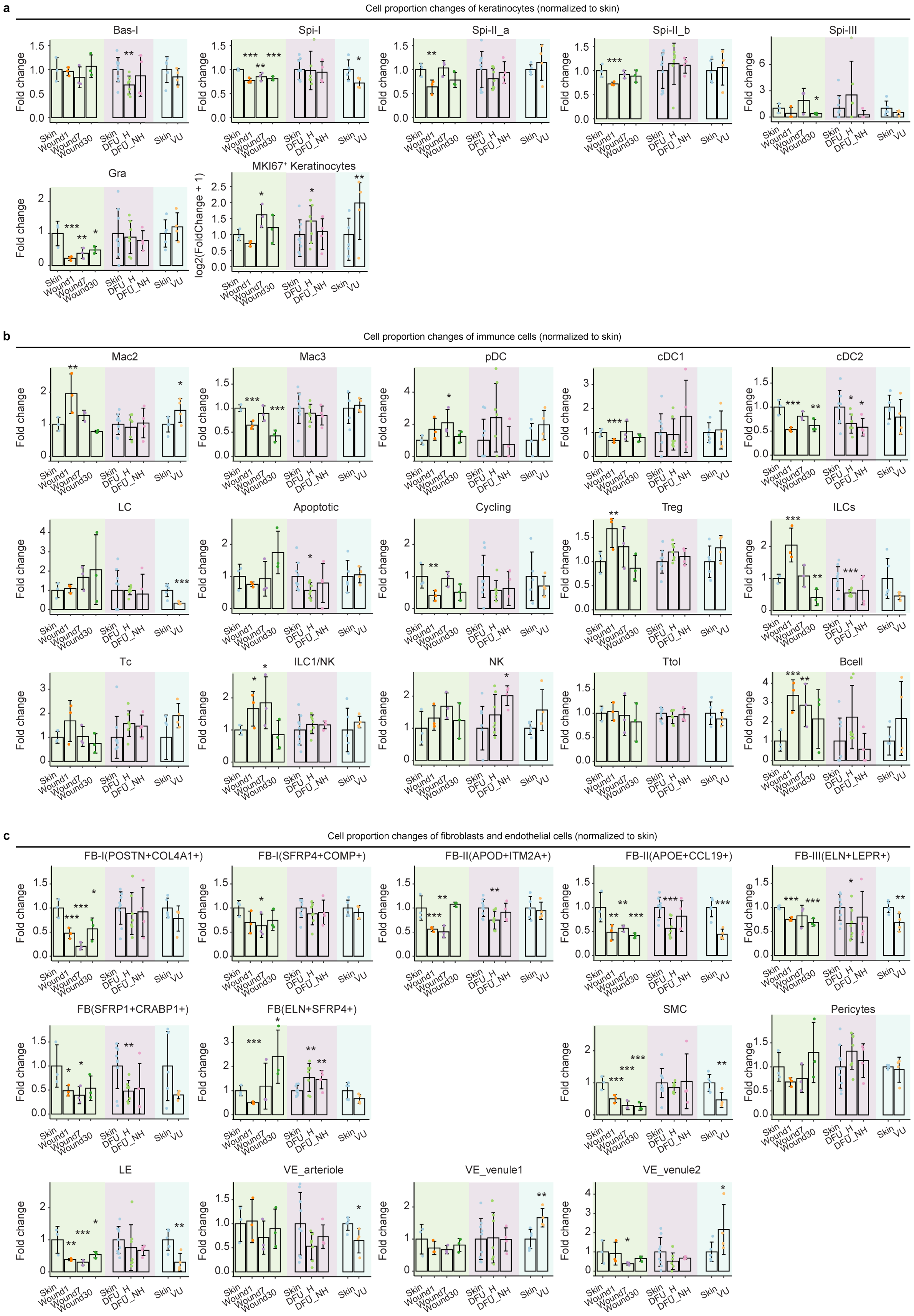

**Figure S7 Multi-facet pathological changes in chronic wounds related to Figure 6.** Bar charts showing fold changes of cell proportions in acute and chronic wounds normalized to healthy skin for keratinocytes (**a**), immune cells (**b**), fibroblasts, and endothelial cells (**c**). Bas-/Spi-/Gra, basal/spinous/granular keratinocyte; Mac2/3: macrophage 2/3; cDC/pDC, conventional/ plasmacytoid dendritic cell; LC, Langerhans cell; Treg, regulatory T cell; ILC, innate lymphoid cell; Tc: cytotoxic T cell; NK-cell, natural killer cell; Ttol, tolerant T cell; FB, fibroblast; SMC, vascular smooth muscle cell; LE/VE, lymphatic/vascular endothelial cell. MKI67 is a marker of proliferating cells. Apoptotic and cycling cells are myeloid cells. Significance was assessed using generalized linear modeling on a quasi-binomial distribution, comparing other conditions to healthy skin, *: $p < 0.05$, **: $p < 0.01$, ***: $p < 0.001$.

**Figure S8 Comparison of human and murine skin wound healing related to Figure 7**

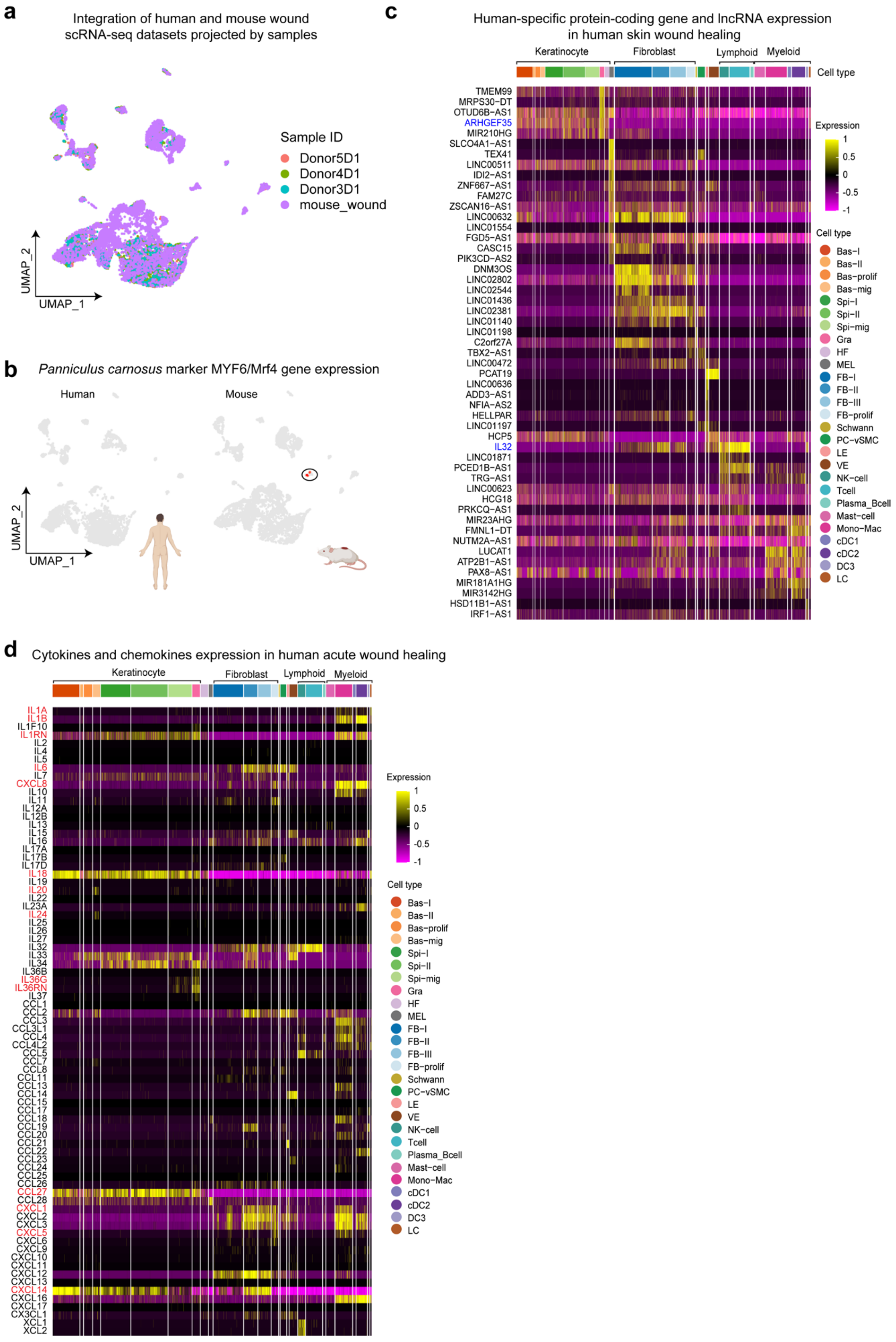

**Figure S8 Comparison of human and murine skin wound healing related to Figure 7.** (**a**) UMAP showing the cell distribution of each sample in integrated scRNA-seq datasets of human and mouse acute wounds after batch correction. (**b**) Feature plots showing marker gene expression (MYF6/Mrf4) of *panniculus carnosus* muscle in integrated human and mouse clusters. Heatmaps showing the expression of human-specific protein-coding and long non-coding RNA genes (**c**), as well as cytokines and chemokines (**d**), in different cell types of human acute wound healing. Genes colored blue in (**c)** are protein-coding genes.

#### Supplementary Schematic summary of the study

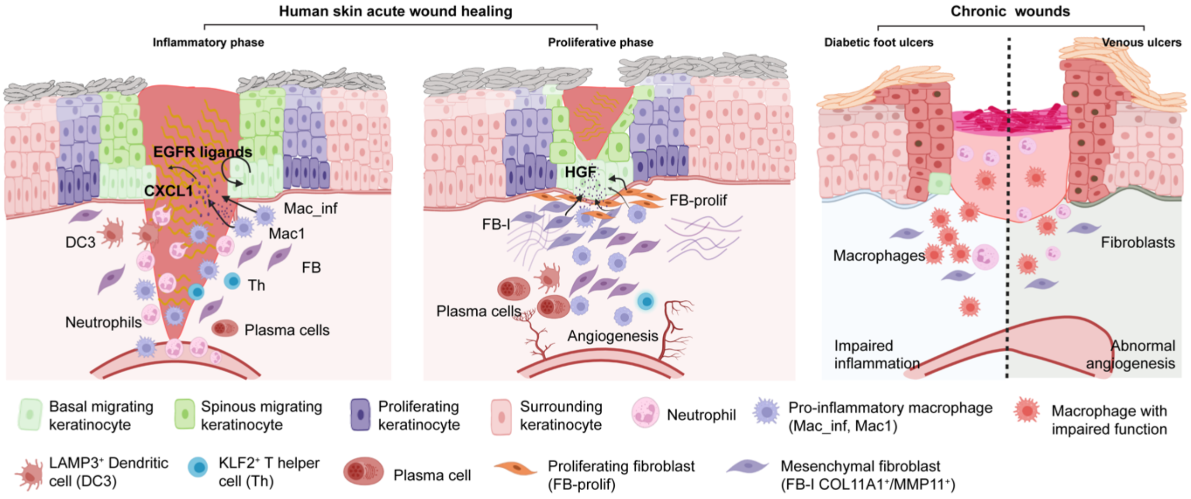
